# RNA-triggered protein cleavage and cell death by the RNA-guided type III-E CRISPR-Cas nuclease-protease complex

**DOI:** 10.1101/2022.08.17.504292

**Authors:** Kazuki Kato, Sae Okazaki, Cian Schmitt-Ulms, Kaiyi Jiang, Wenyuan Zhou, Junichiro Ishikawa, Yukari Isayama, Shungo Adachi, Tomohiro Nishizawa, Kira S. Makarova, Eugene V. Koonin, Omar O. Abudayyeh, Jonathan S. Gootenberg, Hiroshi Nishimasu

## Abstract

The type III-E Cas7-11 effector nuclease forms a complex with a CRISPR RNA (crRNA) and the putative caspase-like protease Csx29, catalyzes crRNA-guided target RNA cleavage, and has been used for RNA targeting in eukaryotic cells. Here, we report cryo-electron microscopy structures of the Cas7-11–crRNA–Csx29 complex with and without target RNA, and demonstrate that target RNA binding induces a conformational change in Csx29 and results in the protease activation. Biochemical analysis confirmed that Cas7-11-bound Csx29 cleaves Csx30 in a target RNA-dependent manner. Reconstitution of the system in bacteria uncovered Csx30-dependent cellular toxicity regulated by Csx31, and that Csx29-mediated cleavage produces toxic Csx30 fragments, promoting cell death. We find that Csx30 can bind both Csx31 and the associated sigma factor RpoE, suggesting Csx30 can inhibit RpoE and modulate cellular stress response towards infection. Overall, the RNA-guided nuclease-protease activities of the Cas7-11–Csx29 effector complex facilitate protease-based programmed cell death.

## Introduction

Prokaryotic CRISPR-Cas systems provide adaptive immunity against foreign nucleic acids, including phages and mobile genetic elements, via diverse mechanisms of programmed nucleic-acid cleavage. CRISPR-Cas systems are divided into two classes based on the number of components in the effector complexes responsible for defense via cleavage of invading nucleic acids programmed by a CRISPR RNA (crRNA) guide(*1*, *2*). In Class 1 systems, which encompass types I, III, and IV, target nucleic acids are degraded by multi-protein effector complexes, whereas, in Class 2 systems, including types II, V, and VI, the effector complexes are formed by a single multidomain Cas protein (Cas9, Cas12, and Cas13, respectively). Beyond primary effector nuclease function, both Class 1 and Class 2 CRISPR-Cas systems deploy a wide-array of accessory proteins to enhance the antiviral activity of the primary effector nuclease, including secondary nuclease activation via cyclic oligoadenylate generation in type III-A/B/D systems(*3*–*6*) and target RNA-dependent pore formation by Csx28 in type VI-B systems(*7*).

Unlike typical Class 1 effectors, the type III-E effector Cas7-11 (also known as gRAMP) is a single-protein, multidomain effector that consists of four Cas7 domains (Cas7.1–Cas7.4) and a Cas11 domain and likely evolved from the more complex type III-D multi-subunit effectors via domain fusions(*8*, *9*). Cas7-11 associates with a crRNA and cleaves complementary single-stranded RNA (ssRNA) targets at two defined positions, using the Cas7.2 and Cas7.3 domains, respectively. Whereas the type VI effector Cas13 displays promiscuous RNase activity(*10*, *11*), Cas7-11 exhibits specific, guide RNA-dependent RNA cleavage activity in human cells, and has been used as a novel RNA-targeting tool with high specificity and low cell toxicity(*8*, *9*). The type III-E locus contains multiple conserved accessory proteins, including Csx29 (a caspase-like putative protease with fused TPR and CHAT domains), Csx30 and Csx31 (proteins with unknown functions), and RpoE (an alternative sigma factor). Cas7-11 associates with Csx29(*8*, *9*), suggesting a potential mechanism of RNA-guided protease activity for antiviral immunity. Recently, we reported the cryo-electron microscopy (cryo-EM) structure of *Desulfonema ishimotonii* Cas7-11 in complex with its cognate crRNA and target RNA (tgRNA)(*12*), providing mechanistic insights into the pre-crRNA processing and tgRNA cleavage. However, how Cas7-11 cooperates with the other proteins encoded in the type III-E locus (Csx29, Csx30, Csx31, and RpoE), and how Cas7-11 binds to Csx29 and potentially activates its protease activity remain unknown.

### Structures of Cas7-11 in complex with Csx29

We reasoned that structural insights would allow for mechanistic understanding of the Cas7-11–Csx29 effector complex. To prepare the Cas7-11–crRNA–Csx29 complex for structural analysis, we co-expressed the catalytically inactive *D. ishimotonii* Cas7-11 mutant (referred to as Cas7-11 for simplicity), with D429A (Cas7.2) and D654A (Cas7.3) mutations introduced to prevent tgRNA cleavage by Cas7-11, together with Csx29 and a crRNA transcribed from a CRISPR array containing two repeat–spacer units. We determined the cryo-EM structures of the Cas7-11–crRNA–Csx29 complex with and without a 27-nucleotide tgRNA at 2.8- and 2.5-Å resolutions, respectively (Fig. 1A–1D, fig. S1A–S1D and S2A–S2D, and Table S1). In the Cas7-11–crRNA–Csx29 complex, Cas7-11 adopts a modular architecture consisting of four Cas7 domains (Cas7.1–Cas7.4) with a zinc finger (ZF) motif, a Cas11 domain, an insertion (INS) domain inserted within the Cas7.4 domain, a C-terminal extension (CTE) domain, and four interdomain linkers (L1–L4) (Fig. 1C and 1D), as in the Csx29-unbound Cas7-11–crRNA–tgRNA structure(*12*) (fig. S3A–S3C).

**Fig. 1.**
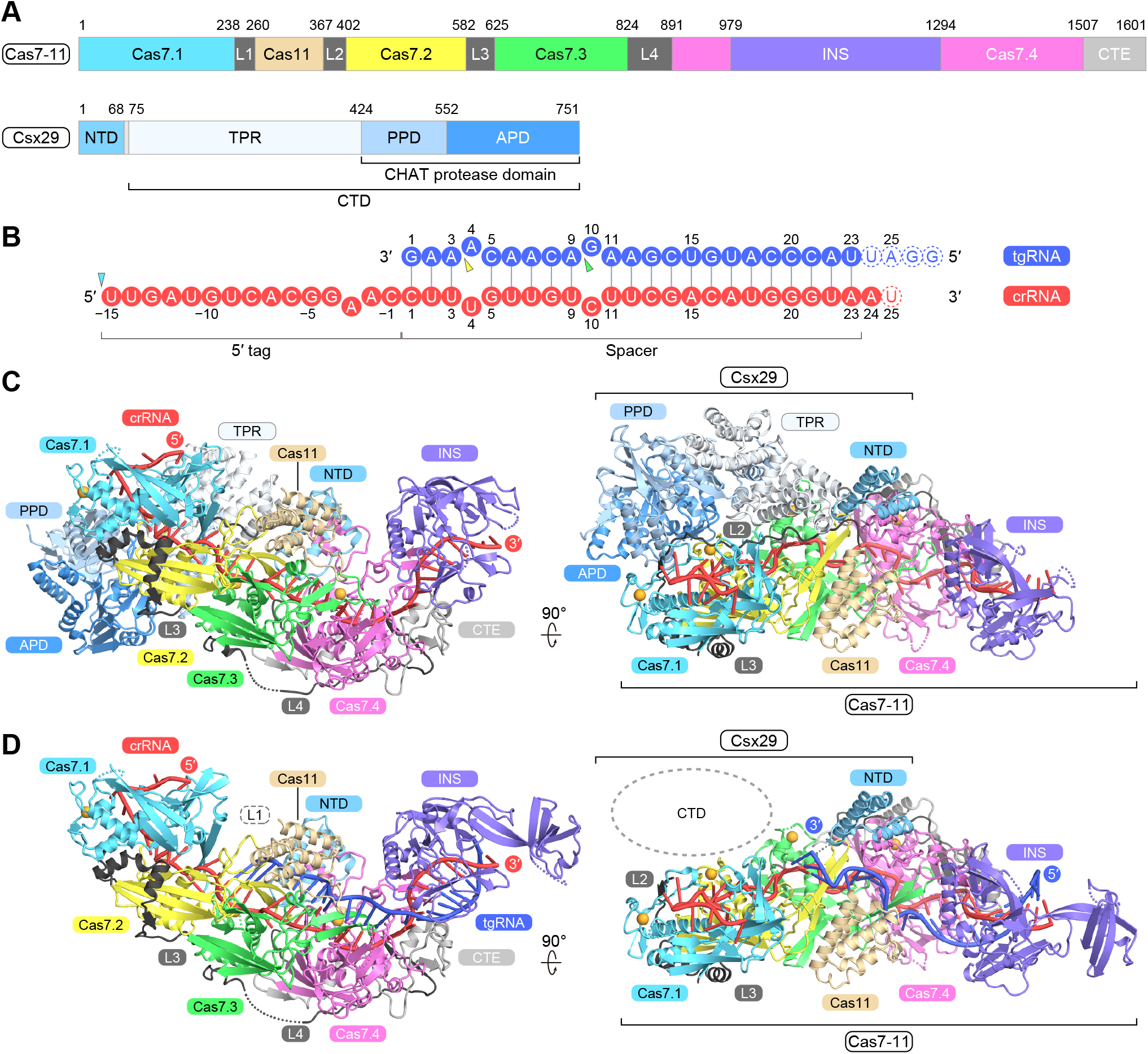
Cryo-EM structures of the Cas7-11–crRNA–Csx29 complexes with and without the target RNA. (A) Domain structures of Cas7-11 and Csx29. (B) Nucleotide sequences of the crRNA and its target RNA. The pre-crRNA processing site and the target RNA cleavage sites are indicated by cyan and yellow/green triangles, respectively. (C and D) Overall structures of Cas7-11–crRNA–Csx29 (C) and Cas7-11–crRNA–Csx29–tgRNA (D). The bound zinc ions are shown as orange spheres. The disordered L1 and L2 linkers are not shown for clarity. The disordered CTD of Csx29 is indicated by a dashed circle in (D).

The 15-nt 5′ tag region (U(−15)–C(−1)) in the 38-nt crRNA (U(−15)–A23) is anchored by the Cas7.1 and Cas7.2 domains (Fig. 1C and 1D). Nucleotide U(−16) was not resolved in the density map (fig. S4A), suggesting that the co-expressed pre-crRNA was processed by Cas7-11 into the mature crRNA. U(−15) is surrounded by H43, R53, Y55, N152, and S158 in the Cas7.1 domain (fig. S4A), consistent with the proposed pre-crRNA processing mechanism, in which H43 functions as a general base to deprotonate the 2′-hydroxy group of U(−16) for pre-crRNA maturation(*12*). In the tgRNA-free structure, the 23-nt crRNA spacer region (C1–A23) is recognized by the Cas7.2–Cas7.4 domains (Fig. 1C and fig. S3B), while in the tgRNA-bound structure, the crRNA spacer region (C1–A23, except for U4 and C10) hybridizes with the tgRNA (G1*–U23*, except for A4* and G10*) to form a guide-target duplex (Fig. 1D and fig. S3C), as in the Cas7-11–crRNA–tgRNA structure(*12*) (fig. S3A). The thumb-like β-hairpins in the Cas7.2 and Cas7.3 domains penetrate the guide-target duplex, inducing base flipping at fourth and tenth positions (U4-A4* and C10-G10*), respectively (fig. S4B). The catalytic residues D429 (Cas7.2) and D654 (Cas7.3) (D429A and D654A in the present structures) are located in the vicinity of the scissile phosphodiester bonds before A4* and G10* of the tgRNA, respectively (fig. S4B), as previously observed(*12*). These observations explain why Csx29 binding does not substantially affect tgRNA cleavage by Cas7-11(*8*, *9*). In the tgRNA-free structure, the peripheral region (residues 1044–1124) of the INS domain was less resolved in the density map (fig. S2A), probably due to its flexibility resulting from the lack of the interaction with the guide-target duplex. Thus, the peripheral region of the INS domain was not included in the final model of the tgRNA-free structure.

### Csx29 structure

Csx29 consists of an N-terminal domain (NTD) (residues 1–67) and a C-terminal domain (CTD) (residues 75–751), connected with a flexible interdomain linker (residues 68–74) that is disordered in the Cas7-11–crRNA–Csx29 structure (Fig. 2A). The NTD adopts a three-helix bundle and interacts with the Cas7.4 domain of Cas7-11 (Fig. 2A). The CTD can be divided into a TRP (tetratricopeptide repeat) domain consisting of eight TPR units (TPR1–TPR8) (residues 75–423) and a CHAT (Caspase HetF Associated with TPRs) protease domain (residues 424–751) (Fig. 2A). In Csx29, each TPR unit contains two α helices, similar to canonical TPR-containing proteins where TPRs interact with their protein targets(*13*). In the Cas7-11–crRNA–Csx29 structure, the TPR domain of Csx29 interacts with the L2 linker of Cas7-11 (Fig. 1C). The CHAT domain of Csx29 consists of a central 11-stranded mixed β-sheet and flanking α-helices, and can be divided into a pseudo-protease domain (residues 424–551) and an active-protease domain (residues 552–751) with the conserved putative catalytic residues H615 and C658 (Fig. 2A). A Dali search(*14*) confirmed that the CHAT domain of Csx29 structurally resembles caspase-like cysteine proteases, such as human separase(*15*) (fig. S5).

**Fig. 2.**
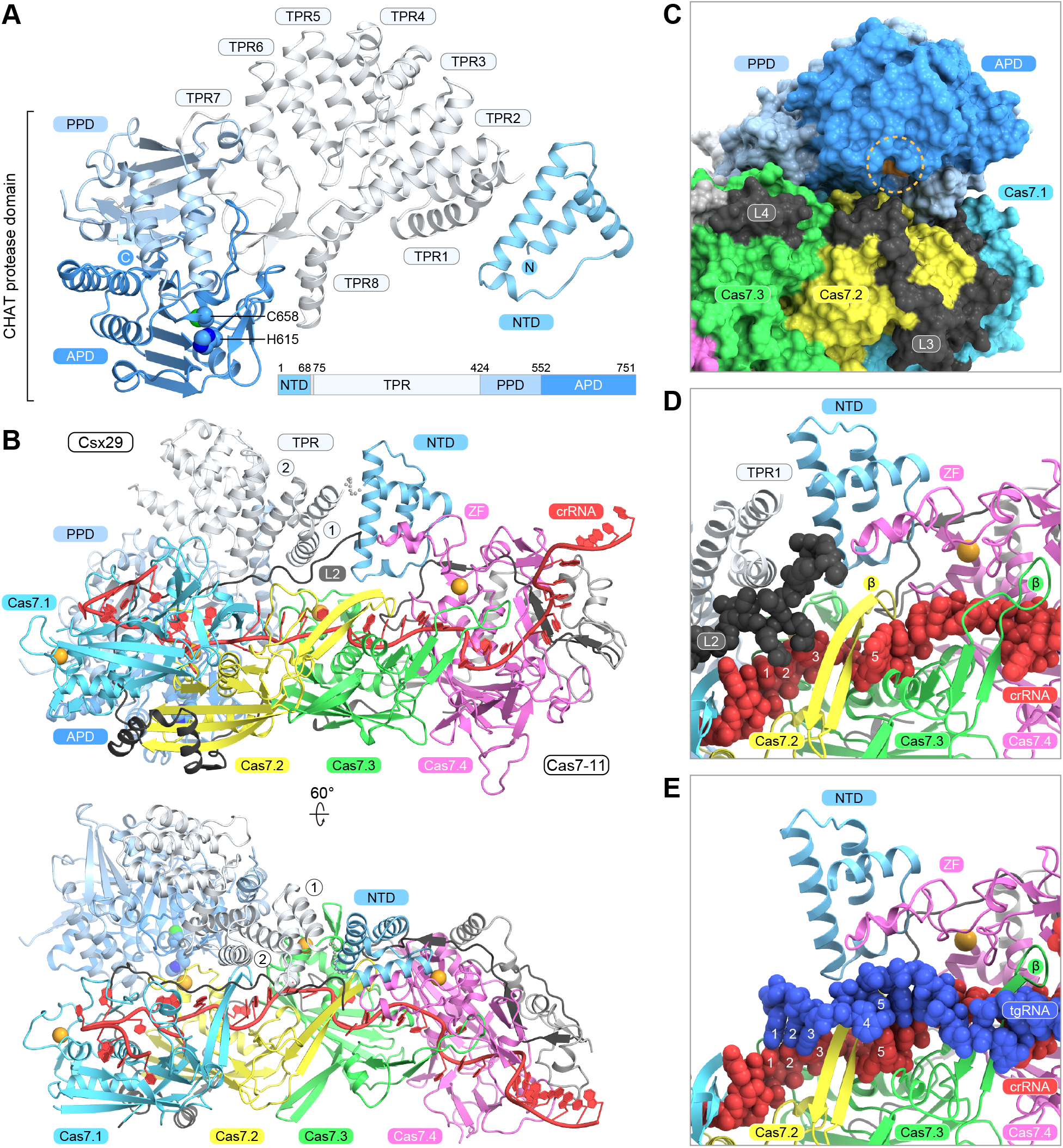
Interaction between Cas7-11 and Csx29. (A) Structure of Csx29 in the Cas7-11–crRNA–Csx29 complex. (B)Interface between Cas7-11 and Csx29 in the Cas7-11–crRNA–Csx29 complex. The Cas11 and INS domains are omitted for clarity. (C) Location of the Csx29 active site. The catalytic residues (H615 and C658) of the Csx29 protease are colored orange. (D and E) Close-up views of the interface between Cas7-11 (Cas7.4 ZF) and Csx29 (NTD). The Cas7-11 L2 linker, crRNA, and tgRNA are shown as space-filling models. The Cas7-11 L2 and Csx29 CTD are disordered in the Cas7-11–crRNA–Csx29–tgRNA structure in (E).

### Interactions between Cas7-11 and Csx29

In the tgRNA-free structure, Csx29 extensively interacts with Cas7-11 at multiple regions (Fig. 2B and fig. S6A). The L2 linker (residues 367–401) and an α-helical insertion (residues 1313–1341) in the Cas7.4 ZF motif, which are disordered in the Csx29-unbound Cas7-11 structure(*12*) (fig. S6B), are ordered and form interactions with Csx29 in the Cas7-11–crRNA–Csx29 structure (Fig. 2B and fig. S6A and S6C). This α-helical insertion in the ZF motif is unique to Cas7.4 and absent in Cas7.1–Cas7.3 (fig. S6C). The NTD of Csx29 mainly interacts with the Cas7.4 domain of Cas7-11 (Fig. 2B and fig. S6A). I5, I8, L30, Y33, L50, R53, F57, L60, S61, and R64 of Csx29 hydrophobically interact with W1316, L1322, L1325, Y1328, L1333, and L1334 of the α-helical insertion region of Cas7-11, while R53 and R64 form hydrogen bonds with R1336 and E1330 of Cas7-11, respectively (fig. S7A). In addition, the Cas7.2 thumb-like β-hairpin and the L2/L4 linkers contribute to the binding to the Csx29 NTD (Fig. 2B). N505 and F507 of Cas7-11 (Cas7.2) interact with T44 and E42/L45 of Csx29, respectively, and K879 and E878 of Cas7-11 (L4) hydrogen bond with E42 and N3/Q47 of Csx29, respectively (fig. S7B). Furthermore, L370 of Cas7-11 (L2) is accommodated within a hydrophobic pocket at the NTD–TPR1 interface of Csx29 (fig. S7C).

TPR1 and TPR2 interact with Cas7.3 (ZF) and L2 of Cas7-11, respectively (Fig. 2B and fig. S6A). R97/E101 of TPR1 and R136 of TPR2 hydrogen bond with D705/Y718 and D705 of Cas7.3 (fig. S7D). TPR2 also interacts with Cas7.1 (thumb-like β-hairpin) and Cas7.2 (ZF) (Fig. 2B). TPR1 and TPR2 are the only TPR domains that mediate the Cas7-11–Csx29 interaction, and TPR3–TPR8 do not contact Cas7-11. The CHAT protease domain of Csx29 interacts with Cas7.1 (thumb-like β-hairpin), Cas7.2 (ZF), Cas7.3 (ZF), and L2 of Cas7-11 (Fig. 2B and fig. S6A). Notably, the protease active site of Csx29 is located in the vicinity of the Cas7.2 domain of Cas7-11 (Fig. 2C), suggesting limited accessibility for the peptide substrate in this conformation. Furthermore, unlike in the separase–securin structure(*15*), the side chain of the catalytic residue C658 is buried inside the CHAT domain in the present structure (fig. S5), indicating that a structural rearrangement of C658 would be required for the substrate cleavage. These observations suggest that the Cas7-11–crRNA–Csx29 structure represents the inactive state of the Csx29 putative protease.

### Target RNA binding-induced structural change in the Cas7-11–Csx29 complex

A comparison of the Cas7-11–crRNA–Csx29 structures with and without the tgRNA revealed a notable conformational difference in Csx29 (Fig. 1C and 1D). While the Csx29 NTD similarly binds Cas7-11 in the two structures (Fig. 1C and 1D), the Csx29 CTD is not resolved in the density map in the tgRNA-bound structure (fig. S2B), indicating that upon the tgRNA binding, the Csx29 CTD dissociates from Cas7-11 and adopts a flexible conformation. This conformational change releases the steric block on the Csx29 active site, allowing access to the substrate protein. A comparison between the two structures indicated that the tgRNA (G1* and A2*) sterically clashes with Cas7-11 (L2) and Csx29 (TPR1) in the Cas7-11–crRNA–Csx29 structure (Fig. 2D, 2E, and fig. S6E), consistent with the tgRNA binding-induced structural change in Csx29. These structural observations suggest that the Cas7-11–crRNA–Csx29–tgRNA structure represents the active state of the Csx29 protease and that Csx29 is a target RNA-triggered protease.

### Target RNA-triggered Csx30 cleavage by Csx29

Given our structural findings, we explored whether proteins encoded in the endogenous *D. ishimotonii* type III-E CRISPR locus are substrates for Csx29 cleavage. Neither Cas7-11 nor Csx29 were degraded in the presence of the tgRNA in biochemical assays (Fig. 3A and 3B), suggesting that the Cas7-11–Csx29 complex itself is not a Csx29 substrate. Given that Csx30 and Csx31 are encoded together with Cas7-11 and Csx29 in the CRISPR operon and are highly conserved among the type III-E systems(*8*), we hypothesized that Csx29 could target either Csx30 or Csx31. To test this hypothesis, we attempted to prepare the recombinant Csx30 and Csx31 proteins and examine whether they are cleaved by Csx29 in a tgRNA-dependent manner. We found that while Csx30 could be purified as a soluble protein, Csx31 was expressed in an insoluble fraction. We examined the *in vitro* cleavage of Csx30 by Cas7-11–crRNA–Csx29 in the absence and presence of the tgRNA, and found that Cas7-11–crRNA–Csx29 cleaves Csx30 into two fragments, Csx30-1 (~50 kDa) and Csx30-2 (~15 kDa), only in the presence of the tgRNA (Fig. 3B). The C615A/H658A mutations in Csx29 abolished the Csx30 cleavage (Fig. 3B), but did not affect the tgRNA cleavage by Cas7-11 (fig. S8), indicating the separable nuclease and protease activities. Furthermore, the D429A/D654A catalytic mutations in Cas7-11 abolished tgRNA cleavage (fig. S8), as previously observed(*12*), and, unexpectedly, improved the Csx30 cleavage by Csx29 to almost complete proteolysis (Fig. 3B). This improvement in the proteolytic activity suggests that the tgRNA dissociates from the effector complex after the Cas7-11-mediated cleavage and that the Csx29 protease is only active as long as a target RNA is bound to the Cas7-11–Csx29 complex. These results demonstrated that Csx30 is cleaved by the CHAT protease domain of Csx29 in a target RNA-dependent manner.

**Fig. 3.**
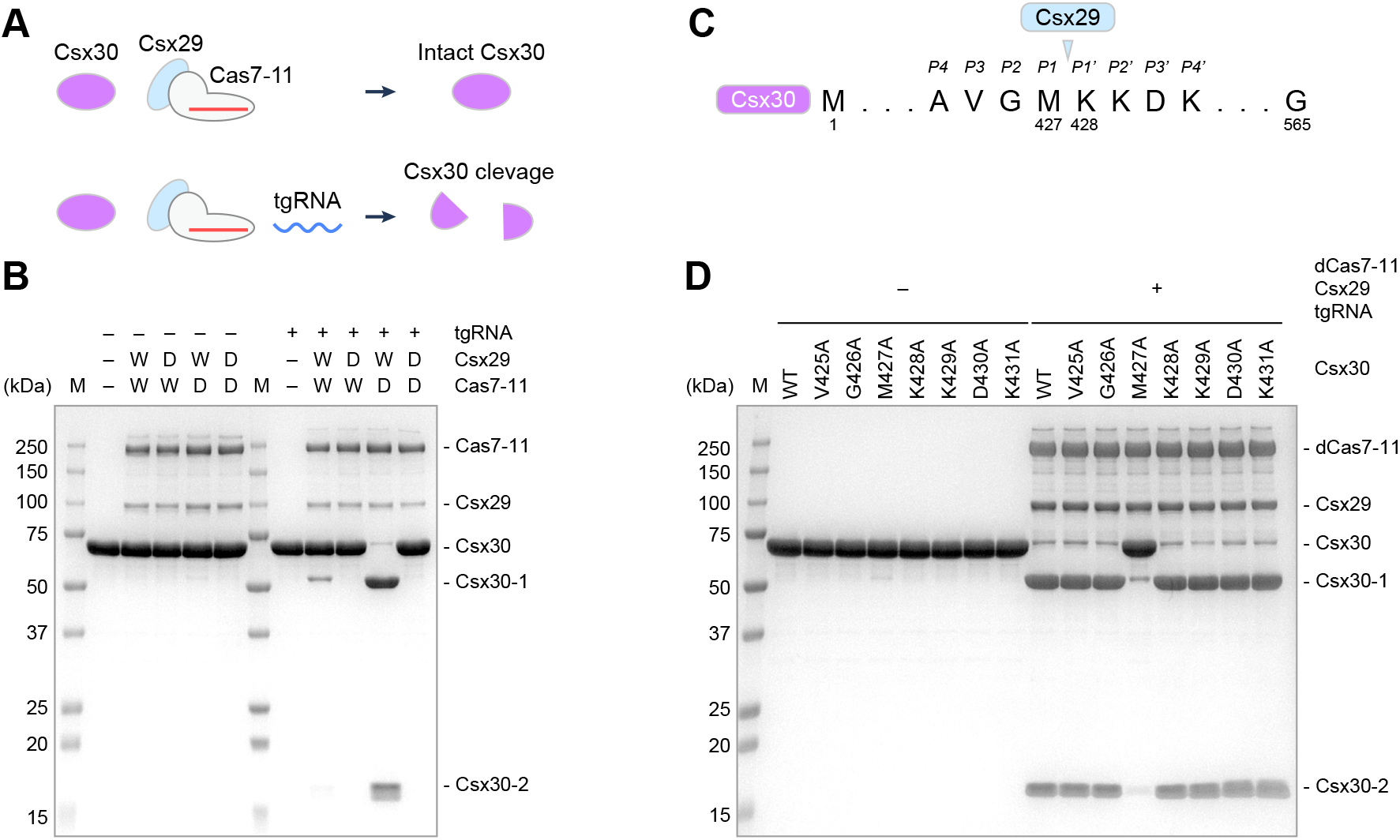
Target RNA-triggered Csx30 cleavage by Csx29. (A) Schematic of the RNA-triggered Csx30 cleavage by the Cas7-11–crRNA–Csx29 complex. (B) RNA-triggered Csx30 cleavage by the Cas7-11–crRNA–Csx29 complex. The Cas7-11–crRNA–Csx29 complex was incubated with Csx30 at 37°C for 2 h in the presence or absence of the target RNA, and then the proteins were analyzed by SDS-PAGE. The wild-type (W) and catalytically inactivated (D) versions of Cas7-11 and Csx29 were used. The gel was stained with CBB. (C) Proteolytic cleavage site in Csx30. The Csx30 site cleaved by Csx29 is indicated by a triangle. (D) Csx29-mediated cleavage of the Csx30 mutants. The dCas7-11–crRNA–Csx29 complex was incubated with the Csx30 mutants at 37°C for 2 h in the presence or absence of the target RNA, and then the proteins were analyzed by SDS-PAGE. The gel was stained with CBB.

N-terminal analysis of the Csx30-2 fragment showed that it begins with K428 (fig. S9), indicating that Csx30 is cleaved by Csx29 between M427 and K428 (Fig. 3C). A structural prediction using AlphaFold2(*16*) indicated that Csx30 consists of an N-terminal domain (NTD) and a C-terminal domain (CTD), which are connected by a linker region. The NTD (residues 1–377) contains two α-helical subdomains, whereas the CTD (residues 418–565) comprises a core β-barrel with flanking α helices (fig. S10). The cleavage site between M427 and K428 is located at a β-hairpin in the Csx30 CTD (fig. S10). We examined the *in vitro* Csx29-mediated cleavage of eight Csx30 mutants, in which V425–K431 were individually replaced with an alanine. The M427A mutation substantially reduced the Csx30 cleavage, whereas the other mutations had no effect (Fig. 3D). Thus, Csx29 seems to primarily recognize M427 at the P1 site within the AVGM|KKDK sequence in Csx30 and cleaves Csx30 between M427 (P1) and K428 (P1′). Together, these results demonstrated that the Cas7-11–crRNA–Csx29 complex catalyzes target RNA-triggered Csx30 proteolytic cleavage.

### Effects of Csx30 and Csx31 on bacterial cell growth

To explore the physiological relevance of the Csx29-mediated Csx30 cleavage, we overexpressed in *Escherichia coli* the full-length Csx30 (referred to as Csx30 for simplicity), the N-terminal fragment of Csx30 (residues 1–427, Csx30-1), or the C-terminal fragment of Csx30 (residues 428–565, Csx30-2), and monitored the cell growth (Fig. 4A). Overexpression of Csx30 substantially inhibited the cell growth compared to uninduced controls (Fig. 4B, 4C, and fig. S11A), demonstrating the cell toxicity of Csx30. Overexpression of Csx30-1 similarly caused significant growth inhibition, whereas Csx30-2 displayed only non-significant, mild inhibition (Fig. 4B, 4C, and fig. S11A), indicating that Csx30-1 harbors the toxicity of this protein. As a structural prediction using AlphaFold2(*16*) suggested that Csx30 and Csx31 have oppositely charged surfaces and electrostatically interact with each other (fig. S11B), we also explored the role of Csx31 in our bacterial growth assays. Overexpression of Csx31 rescued the Csx30-mediated growth defect (Fig. 4D, 4E, and fig. S11C), suggesting that Csx31 interacts with Csx30 and suppresses Csx30-induced toxicity.

**Fig. 4.**
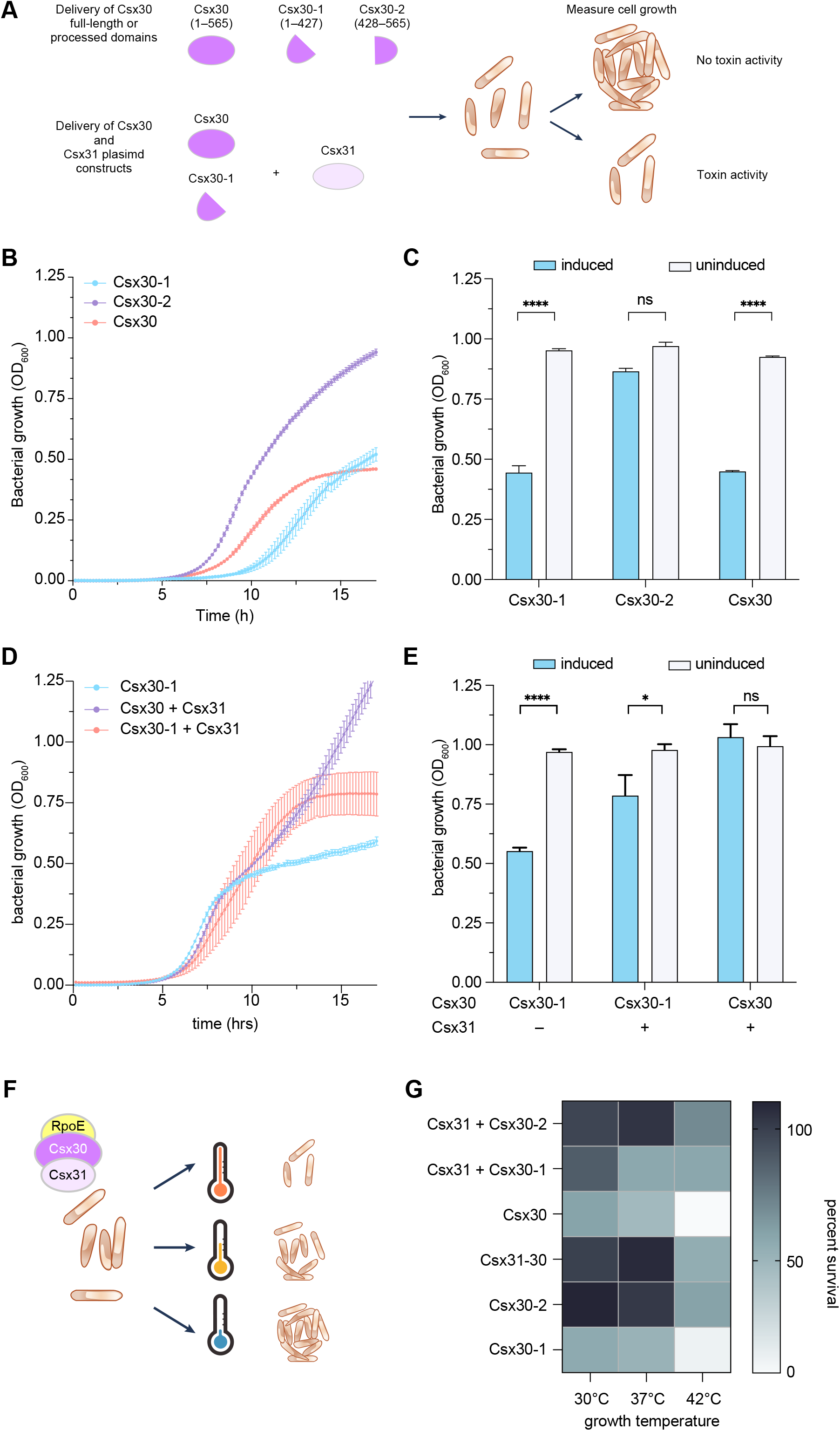
Effects of Csx30 and Csx31 on bacterial cell growth. (A) Schematic of bacterial growth assays for studying the Csx30 and Csx31 functions. (B and C) Growth curves (B) and end-point analysis (C) of *E. coli* expressing either full-length Csx30, the N-terminal fragment (residues 1–427) of Csx30 (Csx30-1), or the C-terminal fragment (residues 428–565) of Csx30 (Csx30-2). (D and E) Growth curves (D) and end-point analysis (E) of *E. coli* expressing either Csx30-1, full-length Csx30 and Csx31, or Csx30-1 and Csx31. In (B)–(E), growth was compared between induced and uninduced expression conditions. In (C) and (E), significance was calculated via two tailed Student’s t test (****, p < 0.0001; n.s., not significant). Data are shown as mean ± s.e.m. (*n* = 3) (F) Schematic of bacterial growth assays at different temperatures ranging from 30°C to 42°C. (G)Heatmap comparing the percent survival of bacteria expressing either Csx30-1, CSX30-2, full length CSX30 and CSX31, full length CSX30, CSX30-1 and CSX31, or CSX30-2 and CSX31 at three different temperatures. (percent survival is calculated by the ratio of OD_600_ of the bacteria in the induced condition over the non-induced condition). Color scale shows percent survival from 0 to 100 percent.

### Interactions between Csx30, Csx31, and RpoE

Csx30, Csx31, and RpoE (a sigma factor involved in the regulation of extracytoplasmic stress response pathways(*17*)) frequently co-occur the type III-E CRISPR loci(*2*, *8*). Based on the results of our bacterial growth assays (Fig. 4), including involvement of Csx31 in modulating Csx30 toxicity, we hypothesized further interplay between the three proteins in the locus. To test this hypothesis, we leveraged the fact that RpoE is involved in the heat shock response in *E. coli* and tested for the effect of Csx30 and Csx31 in *E. coli* at different temperatures from 30°C to 42°C, hypothesizing that the growth defects might be more pronounced at higher temperatures due to inhibition of RpoE by the Csx30 and Csx31 proteins (Fig. 4F). Corroborating our hypothesis, we found that at higher temperatures, the toxicity of Csx30 was more dramatic compared to lower temperatures across all the combinations of proteins tested, suggesting the involvement of RpoE in relation to Csx30 and Csx31 (Fig. 4G). To examine direct interactions between Csx30, Csx31, and RpoE, we co-expressed the three proteins in *E. coli* and analyzed complex formation using gel-filtration. Csx30, Csx31, and RpoE eluted as a single peak from the column (fig. S12A), indicating that they form a stable complex. Like isolated Csx30, Csx30 in the Csx30–Csx31–RpoE complex was cleaved by the Cas7-11–Csx29 complex, and Csx30 (Csx30-1), Csx31, and RpoE co-eluted from the column (fig. S12B), indicating that Csx30-1, Csx31, and RpoE maintain a complex formation after Csx29 cleavage, with separation from Csx30-2. Consistently, structural prediction using AlphaFold2(*16*) suggested that Csx30, Csx31, and RpoE form a ternary complex, in which the Csx30 NTD extensively interacts with RpoE (fig. S12C). *D. ishimotonii* RpoE (DiRpoE) shares structural similarity with *E. coli* RpoE (EcRpoE) (fig. S12D), suggesting that, like the anti-sigma factor RseA(*18*), Csx30 interacts with EcRpoE in our growth assays, causing the cell growth inhibition. While EcRpoE is involved in extracytoplasmic stress response in *E. coli*, the associated regulatory proteins like RseA are not present in *Desulfonema* strains(*19*) and there are different paralogs of RpoE in *E. coli* with varied functions, such as FecI(*20*), suggesting that DiRpoE could mediate an unknown transcriptional response in its host.

A Dali search revealed that the Csx30 CTD is structurally similar to pore-forming proteins in type IV secretion systems, such as CagX(*21*) (fig. S12E). Given its less toxicity in *E. coli* cells, the Csx30 CTD might function as a membrane anchor, rather than a pore-forming protein, consistent with the role of RseA, which is also membrane-localized(*17*, *19*). The CTD and NTD of Csx30 are connected via a flexible linker, suggesting that the Csx29-mediated cleavage releases the N-terminal fragment of Csx30 bound to Csx31 and RpoE into the cytoplasm, thereby modulating gene expression.

Sequence analysis revealed that Csx30 NTDs are highly conserved, whereas Csx30 CTDs are divergent and can be divided into seven distinct groups (fig. S13 and S14), two of which belong to unrelated protein domains found in other contexts. One is an uncharacterized DUF4384 family, which is often fused to different protease domains (see domain architectures for DUF4384 in the CDD database(*22*)). Another group is similar to pilus assembly protein PilP, which forms a periplasmic ring of bacterial type IV pili(*23*). These observations suggest the mechanistic diversity of Csx30-mediated RpoE interaction and programmed gene expression modulation.

## Discussion

In this study, we determined the cryo-EM structures of the Cas7-11–crRNA–Csx29 complex with and without a target RNA. Comparison of the two structures showed that target RNA-binding induces a structural change in the Csx29 protease domain, likely activating the Csx29 protease activity. Consistent with this structural finding, our biochemical analysis demonstrated that Csx29 is a target RNA-triggered protease that cleaves another type III-E associated protein, Csx30, at a unique site. Analysis of the effects of Csx30 and Csx31 on bacterial growth suggested that the Csx29-mediated Csx30 cleavage releases the toxic N-terminal fragment of Csx30 in complex with Csx31 and causes growth inhibition in bacterial cells. Furthermore, our biochemical and structural analyses indicated that Csx30, Csx31, and RpoE can form a ternary complex, in which Csx30 extensively interacts with RpoE, suggesting that Csx30 inhibits the RpoE activity and causes cell growth arrest. Taken together, these findings revealed that the type III-E Cas7-11–Csx29 effector complex is an RNA-triggered programmable nuclease-protease capable of cleaving ssRNA targets and the Csx30 protein, unleashing a downstream signaling cascade that affects cell growth via potential transcriptional regulation.

Thus, in the type III-E CRISPR-Cas systems, the Cas7-11–Csx29 effector complex likely degrades ssRNA transcripts of phage genes and stimulates potentially toxic host cell stress responses through the Csx29-mediated Csx30 cleavage (Fig. 5). Although the exact mechanism of this effect on cell growth remains to be investigated, our finding that Csx30, Csx31, and RpoE form a ternary complex suggests a potential pathway. Conceivably, in the absence of Csx31, the toxic N-terminal fragment of Csx30 might act as an anti-sigma factor, thereby inhibiting the RpoE activity, modulating gene expression, and leading to cell death or growth arrest. Moreover, given unexpected structural similarity between the Csx30 CTD and pore-forming proteins in type IV secretion systems, it is possible that Csx29 proteolysis might liberate the Csx30 NTD from being near the cell membrane into the cytoplasm to exert its cellular effects. This type of programmed cell death might be analogous to that caused by the bacterial membrane pore-forming toxins gasdermins that are activated via proteolytic cleavage and release of auto-inhibitory peptides by associated proteases activated during phage infection(*24*, *25*). Moreover, given the high diversity of Csx30 CTDs (fig. S14), additional exploration of other type III-E systems might reveal other functions associated with Cas7-11-mediated target RNA recognition.

**Fig. 5.**
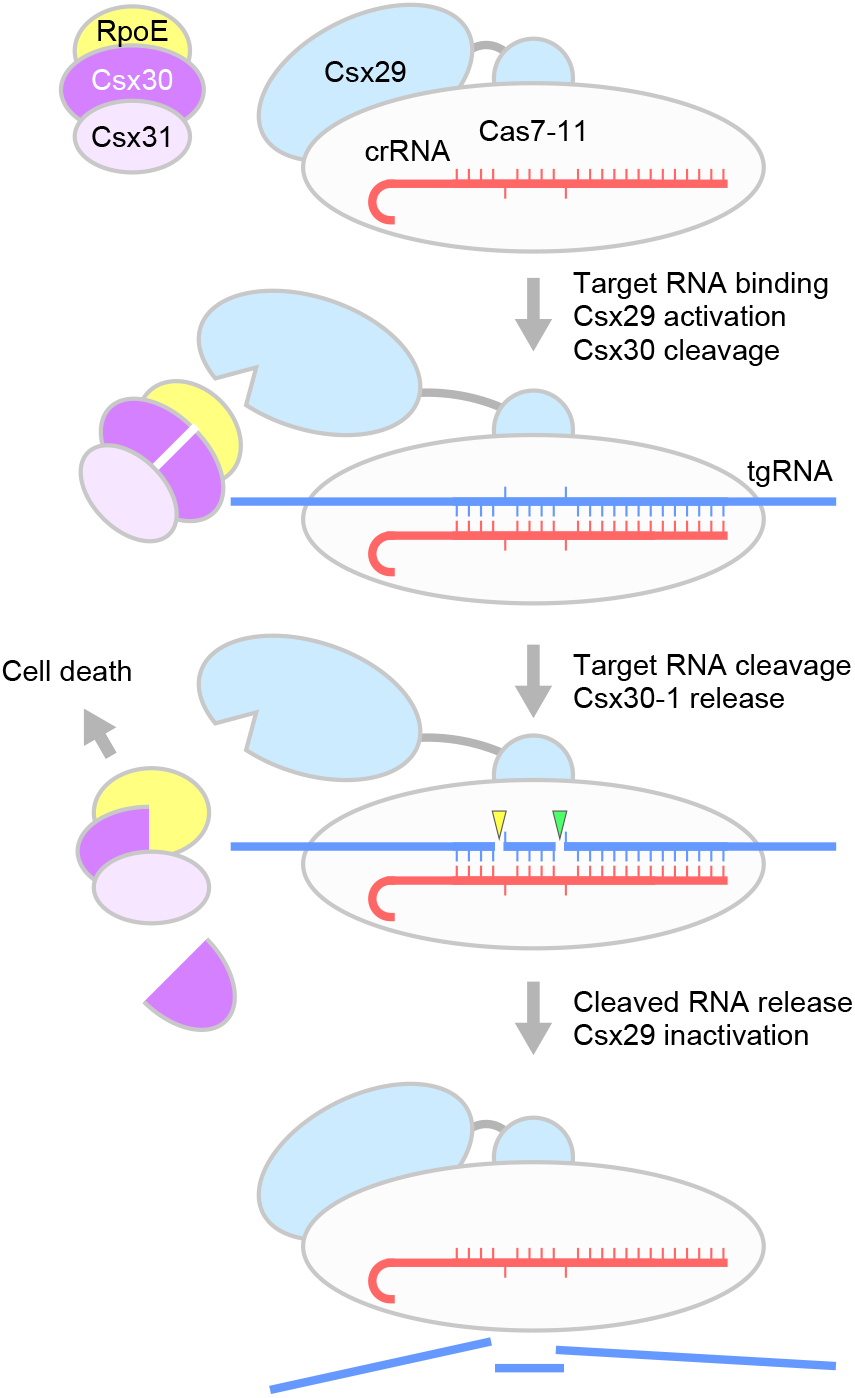
Potential mechanism of cell growth inhibition by the Cas7-11–Csx29 effector complex. The schematic presents a proposed mechanism of the RNA-triggered proteolytic activation of Csx30 by the Cas7-11–Csx29 complex, which induces cell death as part of anti-viral immunity. Csx31 likely functions as an antitoxin, thereby protecting the cell from the toxic effect of the Csx30 NTD. The Csx30 NTD bound to RpoE could affect cell growth and viability through unknown mechanisms.

Among the CRISPR-Cas systems, a biological if not mechanistic analogy can be found in the type VI system, where the Cas13–crRNA effector complex recognizes complementary phage mRNAs and cleaves both phage (specifically and in *cis*) and host (indiscriminately and in *trans*) transcripts, stalling the cell growth and with it, the infectious cycle(*26*). Similarly, within type III systems, the CRISPR-Lon protease can be activated via cyclic oligoadenylates upon RNA recognition by type III effector complexes, activating nearby protein toxins via proteolytic cleavage(*27*). Our characterization of the type III-E system highlighted the remarkable diversity of CRISPR-associated functions activated by programmable nucleic-acid recognition, thereby motivating continued exploration of CRISPR-associated proteins and their potential programmable functions that may have useful roles for biology applications. Our findings that the type III-E Cas7-11–Csx29 effector complex is a so far unique RNA-triggered nuclease-protease establish a new paradigm of prokaryotic signal transduction, and could pave the way for the development of new RNA/protein-targeting technologies, including *in vitro* diagnostics and cellular RNA sensing.

## Acknowledgements

We thank the staff scientists at The University of Tokyo’s cryo-EM facility, especially Y. Sakamaki, for help with cryo-EM data collection, and Dr. Nobuaki Okumura for help with N-terminal analysis. H.N. is supported by the Platform Project for Supporting Drug Discovery and Life Science Research (Basis for Supporting Innovative Drug Discovery and Life Science Research (BINDS)) from AMED under grant number JP21am0101115 (support number 2792); AMED under grant number JP21wm0325048h0001; JSPS KAKENHI Grant Numbers 20K20579 and 21H05281; Takeda Medical Research Foundation; and the Inamori Research Institute for Science. J.S.G. and O.O.A. are supported by NIH grants 1R21-AI149694, R01-EB031957, and R56-HG011857; The McGovern Institute Neurotechnology (MINT) program; the K. Lisa Yang and Hock E. Tan Center for Molecular Therapeutics in Neuroscience; the G. Harold & Leila Y. Mathers Charitable Foundation; MIT John W. Jarve (1978) Seed Fund for Science Innovation; FastGrants; the Cystic Fibrosis Foundation; Google Ventures; Longevity Impetus Grant from Norn Group; NHGRI/TDCC Opportunity Fund; and the McGovern Institute.

## Author contributions

K.K. and S.O. performed sample preparation and biochemical analysis with assistance from Y.I. and J.I.; K.K., S.O., and H.N. performed structural analysis with assistance from J.I. and T.N.; C.S., K.J., W.Z., J.S.G., and O.O.A. performed bacterial experiments; S.A. performed N-terminal analysis; K.S.M. and E.V.K. performed computational analysis; J.S.G., O.O.A., and H.N. wrote the manuscript with help from K.K. and E.V.K; J.S.G., O.O.A., and H.N. conceived the project and supervised the research.

## Data and code availability

The EM densities have been deposited in the Electron Microscopy Public Image Archive under the accession codes 33695 and 33696. The model coordinates have been deposited in the Protein Data Bank under the accession codes 7Y7X and 7Y7Y.

## Method details

### Plasmid construction

For the bacterial expression of the *D. ishimotonii* Cas7-11–crRNA–Csx29 complex, the gene encoding Cas7-11 was amplified by PCR and cloned into the modified pACYCDuet-1 plasmid vector (Novagen), expressing Cas7-11 with an N-terminal maltose-binding protein (MBP) and a C-terminal His_6_-tag (MBP–Cas7-11–His_6_). The gene encoding Csx29 was cloned into the His_6_-Twin-Strep-SUMO-pET28a vector, expressing Csx29 with an N-terminal His_6_–Twin-Strep–SUMO tag (His_6_–Twin-Strep–SUMO–Csx29). The CRISPR array containing two direct repeats interspaced by a spacer with the 5′ LacI-repressed T7 promoter and 3′ T7 terminator sequences was synthesized by Eurofins Genomics. For the bacterial expression of Csx30 and Csx30–Csx31–RpoE, the gene encoding Csx30 or Csx30 and Csx31 was amplified from the type III-E *D. ishimotonii* CRISPR locus and cloned into the modified pE-SUMO vector (LifeSensors), in which the SUMO-coding region is replaced with the HRV3C protease recognition site. The gene encoding RpoE was cloned into the pACYCDuet-1 vector, expressing RpoE with an N-terminal His_6_-tag. The mutants of Cas7-11, Csx29, and Csx30 were generated by a PCR-based method, and the sequences were confirmed by DNA sequencing.

### Sample preparation

Cas7-11, Csx29, and the CRISPR array were co-expressed in *E. coli* BL21 (DE3) (Novagen) by induction with 0.25 mM isopropyl β-D-thiogalactopyranoside (Nacalai Tesque) at 18°C overnight. The *E. coli* cells were lysed by sonication in buffer A (20 mM Tris-HCl, pH 7.5, 20 mM imidazole, 150 mM NaCl, 10% glycerol, and 3 mM 2-mercaptoethanol), and the lysate was clarified by centrifugation at 40,000 g. The supernatant was applied to Ni-NTA Superflow resin (QIAGEN), and the bound protein was eluted with buffer A containing 300 mM imidazole. The eluted fraction was diluted 2-fold with buffer B (20 mM Tris-HCl, pH 7.5, 150 mM NaCl, 10% glycerol, and 3 mM 2-mercaptoethanol), and applied to Strep-Tactin XT high capacity (IBA), and the bound protein was eluted with buffer C (100 mM Tris-HCl, pH 8.0, 150 mM NaCl, 10% glycerol, 1 mM EDTA, 50 mM biotin, and 3 mM 2-mercaptoethanol). The eluted protein was applied to Amylose resin (NEB) equilibrated with buffer D (20 mM Tris-HCl, pH 7.5, 150 mM NaCl, 10% glycerol, and 1 mM DTT). The resin was washed with buffer D (20 mM Tris-HCl, pH 7.5, 150 mM NaCl, 10% glycerol, and 1 mM DTT), and the bound protein was eluted with buffer D containing 10 mM D-maltose. For biochemical experiments, the eluted protein was dialyzed against buffer D to remove D-maltose. The concentration of the Cas7-11–Csx29–crRNA complex was measured using Pierce 660-nm Protein Assay Reagent (Thermo Fisher Scientific). Csx30 was expressed in *E. coli* Rosetta2 (DE3), and purified by Ni-NTA Superflow resin and a Superdex 200 Increase column (GE Healthcare). Csx30, Csx31, and RpoE were co-expressed in *E. coli* BL21 (DE3), and the Csx30–Csx31–RpoE complex was purified by Ni-NTA Superflow resin and a HiLoad 16/600 Superdex 200 column (GE Healthcare). The target RNA (25 nucleotides plus 5′ GG) was transcribed *in vitro* with T7 RNA polymerase, and purified by 10% denaturing (7 M urea) polyacrylamide gel electrophoresis. The purified materials were stored at −80°C until use.

### Cryo-EM grid preparation and data collection

To prevent tgRNA cleavage, the catalytically inactive Cas7-11 (D429A/D654A) was used for cryo-EM studies. The Cas7-11–crRNA–Csx29–tgRNA complex was reconstituted by mixing the purified Cas7-11–crRNA–Csx29 complex and the target RNA, at a molar ratio of 1:4. The complex was purified by size-exclusion chromatography on a Superose 6 Increase 10/300 column (GE Healthcare), equilibrated with buffer E (20 mM HEPES-NaOH, pH 7.0, 150 mM NaCl, 1 mM MgCl_2_, and 1 mM DTT). The peak fraction containing Cas7-11–crRNA–Csx29–tgRNA was analyzed by TBE-urea gel, and concentrated to an A_260_ of 3.0, using an Amicon Ultra-4 Centrifugal Filter Unit (MWCO 50 kDa). The sample (3 μL) was applied to freshly glow-discharged Au 300 mesh R1.2/1.3 grids (Quantifoil) in a Vitrobot Mark IV (FEI) at 4°C, with a waiting time of 10 s and a blotting time of 4 s under 100% humidity conditions. The grids were plunge-frozen into liquid ethane cooled at liquid nitrogen temperature. For the cryo-EM analysis of the Cas7-11–crRNA–Csx29 complex, the purified complex was further polished by size-exclusion chromatography on a Superose 6 Increase 10/300 column, equilibrated with buffer E. The peak fraction containing Cas7-11–crRNA–Csx29 was concentrated to an A_260_ of 2.5. The sample was applied onto freshly glow-discharged Au 300 mesh R0.6/1 grids (Quantifoil) in a Vitrobot Mark IV at 4°C, with a waiting time of 10 s and a blotting time of 4 s under 100% humidity conditions. The cryo-EM data were collected using a Titan Krios G3i microscope (Thermo Fisher Scientific), running at 300 kV and equipped with a Gatan Quantum-LS Energy Filter (GIF) and a Gatan K3 Summit direct electron detector. Micrographs were recorded at a nominal magnification of ×105,000 with a pixel size of 0.83 Å in a total exposure of 48 e^−^/Å^2^ per 48 frames. The data were automatically acquired by the image shift method using the EPU software (Thermo Fisher Scientific), with a defocus range of −0.8 to −2.0 μm, and 2,924 and 3,179 movies were acquired for Cas7-11–crRNA–Csx29 and Cas7-11–crRNA–Csx29–tgRNA, respectively.

### Image processing

The data processing was performed using cryoSPARC v3.3.1 software packages(*29*). The dose-fractionated movies were aligned using the Patch motion correction and the contrast transfer function (CTF) parameters were estimated using Patch-Based CTF estimation. Particles were automatically picked using Blob picker and template picker followed by reference free 2D classification to curate particle sets. The particles were further curated by Heterogeneous Refinement using the map derived from cryoSPARC *Ab initio* Reconstruction as a template. The selected particles after Heterogeneous Refinement were refined using Non-uniform refinement(*30*). Local motion correction followed by Non-uniform refinement with optimization of CTF value yielded a map at 2.48 Å and 2.77 Å resolution for Cas7-11–crRNA–Csx29 and Cas7-11–crRNA–Csx29–tgRNA, respectively, according to the Fourier shell correlation (FSC) = 0.143 criterion(*31*). The local resolution was estimated by BlocRes in cryoSPARC.

### Model building and validation

For model building for Cas7-11, the previously published Cas7-11 structure (PDB ID: 7WAH) was rigid-body fitted into the reconstructed density maps in UCSF ChimeraX(*32*). The initial model of Cs×29 was predicted using AlphaFold2(*16*), and fitted into the density map using the Dock Predicted Model in PHENIX package(*33*). These models were manually modified using COOT(*34*) against the density map sharpened using DeepEMhancer(*35*). The models were refined using Real-space refinement in PHENIX(*36*) with the secondary structure and the Ramachandran restraints. Since the MBP, His_6_ and SUMO tags were not resolved in the density map, they were not included in the final models. The structures were validated using MolProbity(*37*) from the PHENIX package. The curve representing model vs. full map was calculated using phenix.mtriage(*38*), based on the final model and the full, filtered and sharpened map. The statistics of the 3D reconstruction and model refinement are summarized in Table S1. The cryo-EM density maps were calculated with UCSF ChimeraX, and molecular graphics figures were prepared with CueMol (http://www.cuemol.org).

### *In vitro* Csx30 cleavage experiment

The purified Cas7-11–crRNA–Csx29 complex (750 nM) was incubated at 37°C for 2 h with the purified Csx30 protein (15 μM) in the presence or absence of the tgRNA (3 μM) in reaction buffer (20 mM HEPES-NaOH, pH 7.5, 150 mM NaCl, 5 mM MgCl_2_, and 2 mM DTT). The reaction was quenched by the addition of an SDS-PAGE sample buffer, and the mixture was then analyzed by SDS-PAGE. The gels were stained with Bullet CBB Stain One (Nacalai Tesque), and then imaged using a FUSION Solo S system (Vilber Bio Imaging).

### *In vitro* target RNA cleavage experiment

The purified Cas7-11–crRNA–Csx29 complex (200 nM) was incubated at 37°C for 10 min with a 5′-Cy5-labeled ssRNA target (600 nM) in reaction buffer (20 mM HEPES-NaOH, pH 7.5, 50 mM NaCl, 5 mM MgCl_2_, and 2 mM DTT). The reaction was quenched by the addition of quenching solution (0.45 mg/mL proteinase K (Nacalai Tesque), 6 mM EDTA, and 200 μM urea), and then incubated at 50°C for 15 min. The mixture was incubated at 100°C for 2 min with 4.5 M urea denaturing buffer, and then analyzed using a 15% Novex PAGE Tris–borate–EDTA (TBE)–urea gel (Invitrogen). The gels were imaged using a FUSION Solo S system, using either Cy5 fluorescence or SYBR Gold fluorescence (Thermo Fisher Scientific).

### N-terminal analysis

The purified nuclease-inactivated Cas7-11–crRNA–Csx29 complex (3 μM) was incubated at 37°C for 1.5 h with the purified Csx30 protein (15 μM) in the presence of the tgRNA (12 μM) in reaction buffer (20 mM HEPES-NaOH, pH 7.5, 150 mM NaCl, and 2 mM DTT). The proteins were separated by SDS-PAGE, blotted onto a PVDF membrane, and stained with Bullet CBB Stain One. The protein band corresponding to a ~15 kDa Csx30 fragment (Csx30-2) was cut out from the membrane, and its N-terminal amino-acid sequence was analyzed by Edman microsequencing(*39*) on a Procise 494 cLC protein sequencer (Applied Biosystems), using the standard pulsed-liquid program for PVDF-blotted proteins.

### Bacterial cell growth experiment

*E. coli* DH5α Competent Cells (Thermo Fisher Scientific) were transformed with a Csx30-expressing plasmid alone or plasmids expressing both Csx30 and Csx31 (Table S2). Single colonies were picked into Terrific broth (Fisher Scientific) containing relevant antibiotics and 1% v/v glucose, and then cultured at 37°C overnight. Following overnight culture, bacterial OD_600_ values were measured and normalized to an initial OD_600_ of 0.00046 in 200 μL final volume by dilution in Terrific broth containing relevant antibiotics as well as 1% glucose for non-induced conditions and 1% arabinose for induced conditions. Using BioCoat Cellware 96-well tissue culture plates (Corning), growth assays were performed using a BioTek Synergy Neo2 at 37°C with continuous shaking, reading OD_600_ values at 10 min intervals for up to 22 hours. For growth curves with different temperatures, the same culturing conditions were used and the BioTek Synergy Neo2 is set to either 30°C, 37°C, or 42°C with continuous shaking for up to 22 hours during the measurement.

### *In vitro* binding experiment

The purified Csx30–Csx31–RpoE complex (15 μM) was incubated at 37°C for 2 h with the purified nuclease-inactivated Cas7-11–crRNA–Csx29 complex (750 nM) in the presence of the tgRNA (3 μM) in reaction buffer (20 mM HEPES-NaOH, pH 7.5, 150 mM NaCl, 5 mM MgCl_2_, and 2 mM DTT). The mixture was separated on a Superdex 200 Increase 10/300 column equilibrated with buffer (20 mM HEPES-NaOH, pH 7.5, 150 mM NaCl, and 2 mM DTT). The peak fractions were analyzed by SDS-PAGE. The gels were stained with Bullet CBB Stain One, and then imaged using a FUSION Solo S system.

---

**fig. S1.**
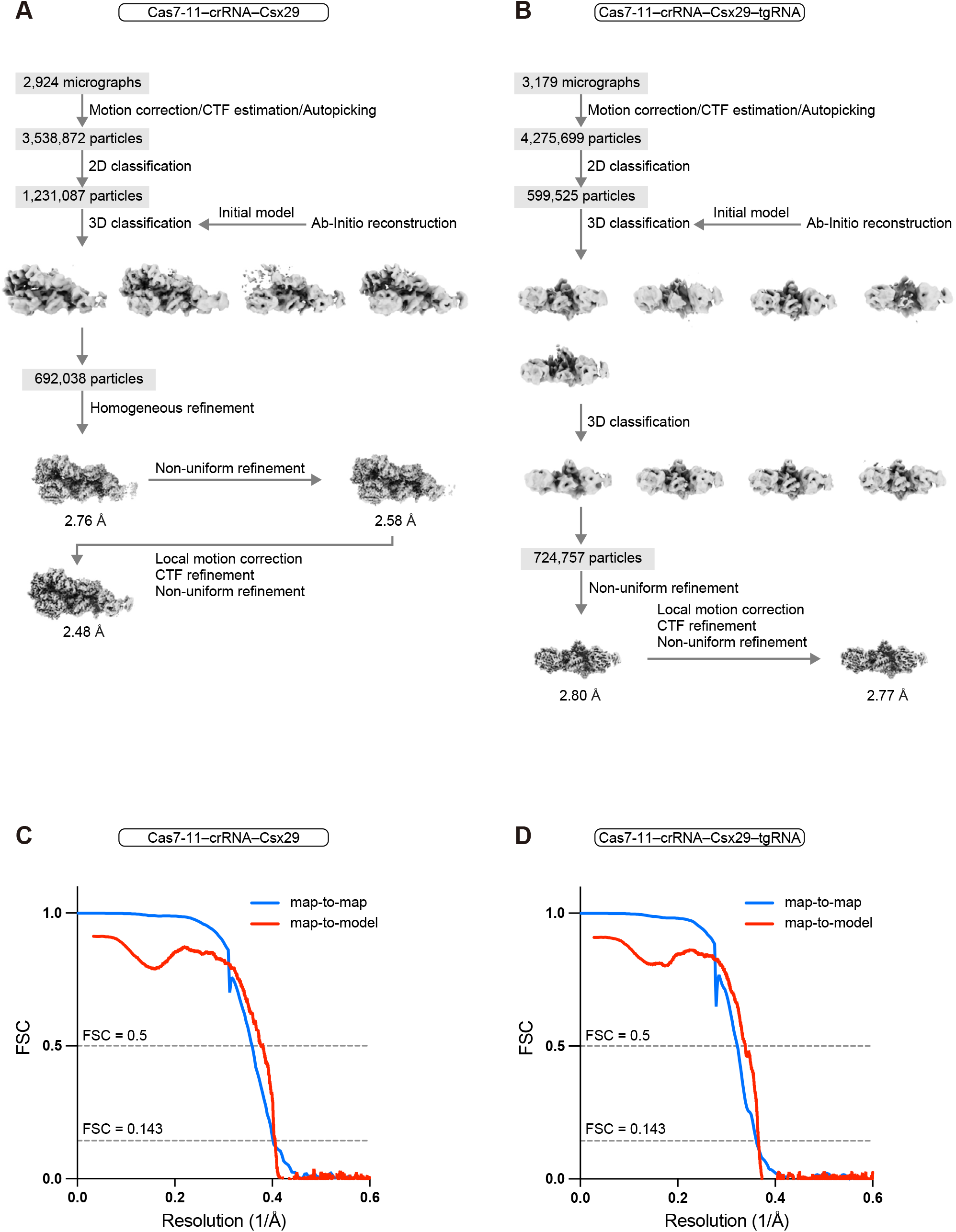
Cryo-EM analysis. (A and B) Single-particle cryo-EM image processing workflows for Cas7-11–crRNA–Csx29 (A) and Cas7-11–crRNA–Csx29–tgRNA (B). (C and D) Fourier shell correlation curves calculated between the half-maps and between the refined model and the density map for Cas7-11–crRNA–Csx29 (C) and Cas7-11–crRNA–Csx29–tgRNA (D).

**fig. S2.**
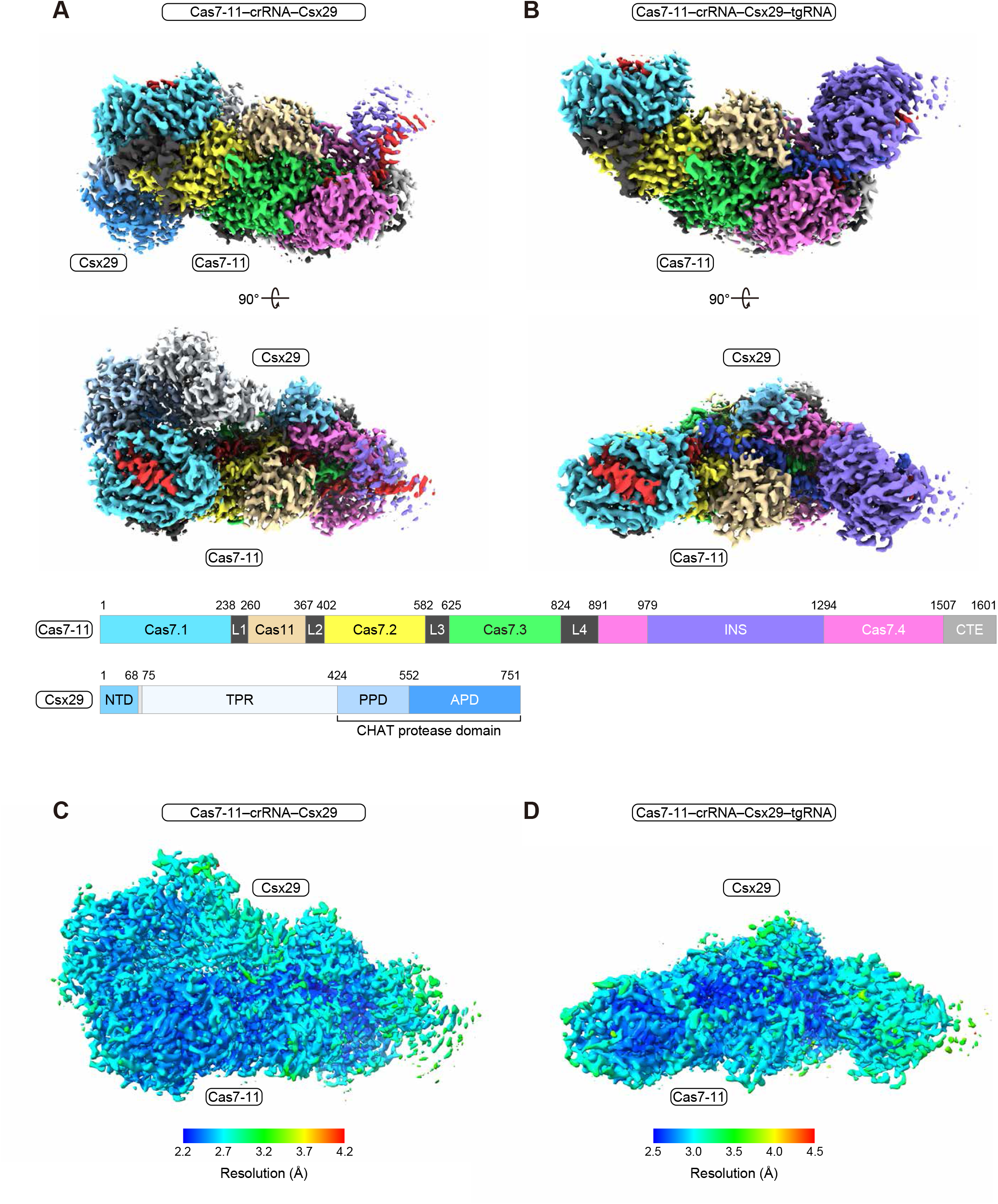
Cryo-EM density maps. (A and B) Cryo-EM density maps, colored according to the protein domains, for Cas7-11–crRNA–Csx29 (A) and Cas7-11–crRNA–Csx29–tgRNA (B). (C and D) Cryo-EM density maps, colored according to the local resolution, for Cas7-11–crRNA–Csx29 (C) and Cas7-11–crRNA–Csx29–tgRNA (D).

**fig. S3.**
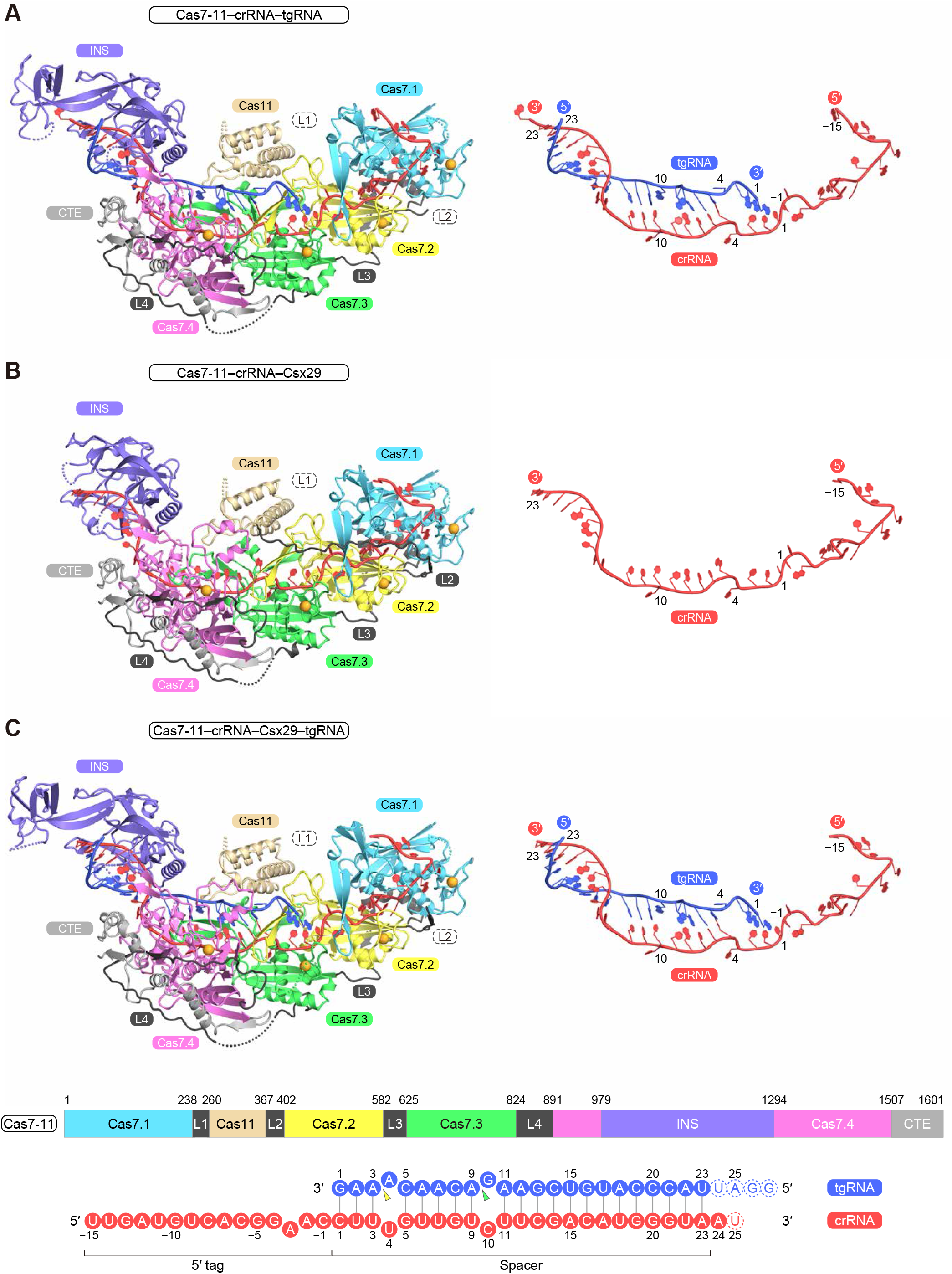
Structural comparison of the Cas7-11 complexes in different states. (A–C) Structures of Cas7-11–crRNA–tgRNA (PDB ID: 7WAH) (A), Cas7-11–crRNA–Csx29 (B), and Cas7-11–crRNA–Csx29–tgRNA (C). The bound zinc ions are shown as orange spheres. The disordered L1 and L2 linkers are not shown for clarity. The bound RNA molecules are shown on the right of the complexes.

**fig. S4.**
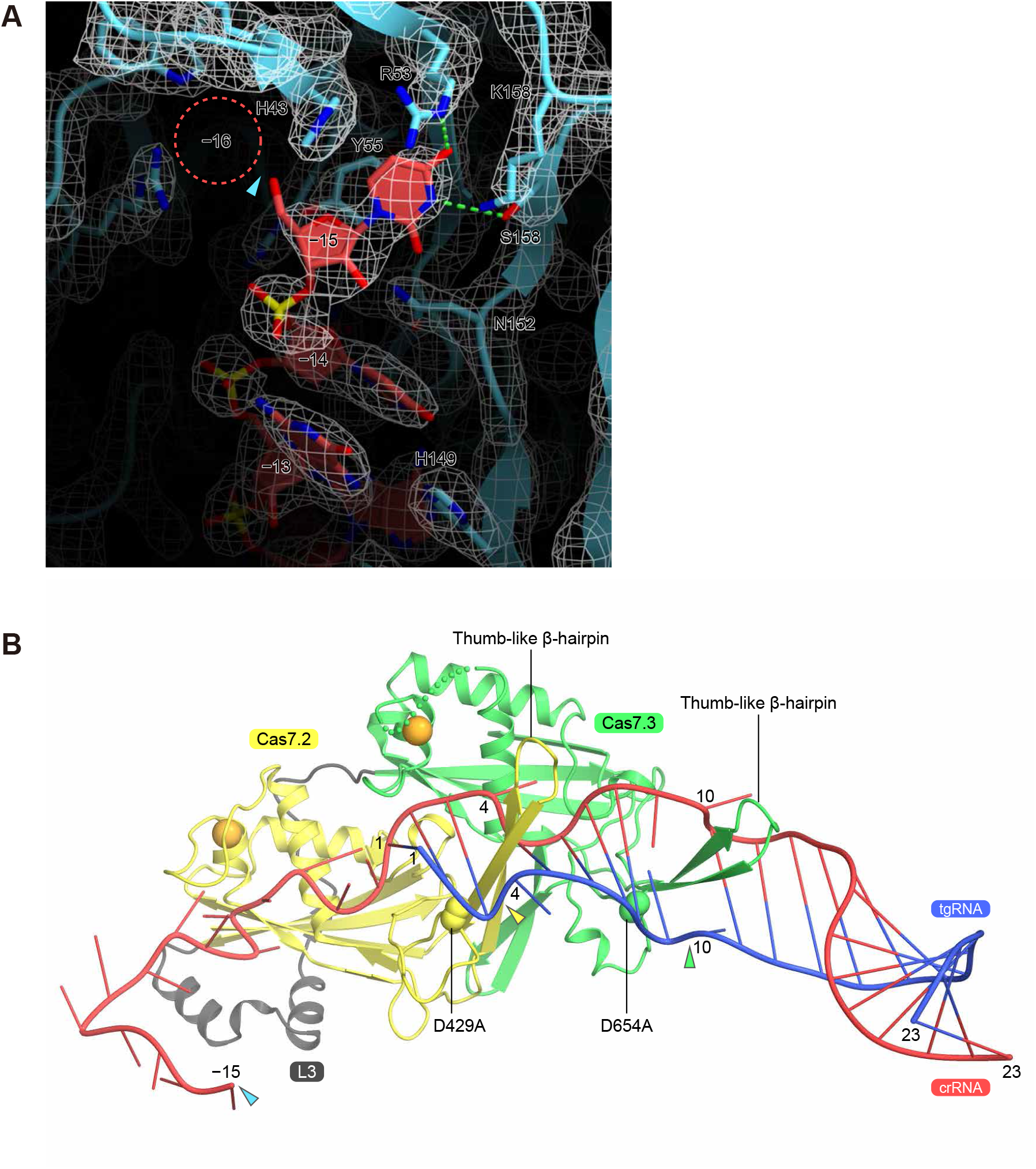
RNA recognition by Cas7-11. (A) Recognition of the crRNA 5′ end by the Cas7.1 domain. The density map is shown as a gray mesh. The possible location of U(−16) and the pre-crRNA processing site are indicated by a dashed circle and a cyan triangle, respectively. (B) Recognition of the guide-target duplex by the Cas7.2 and Cas7.3 domains. The catalytic residues (D429A and D654A) are depicted as space-filling models. The target RNA cleavage sites are indicated by yellow and green triangles.

**fig. S5.**
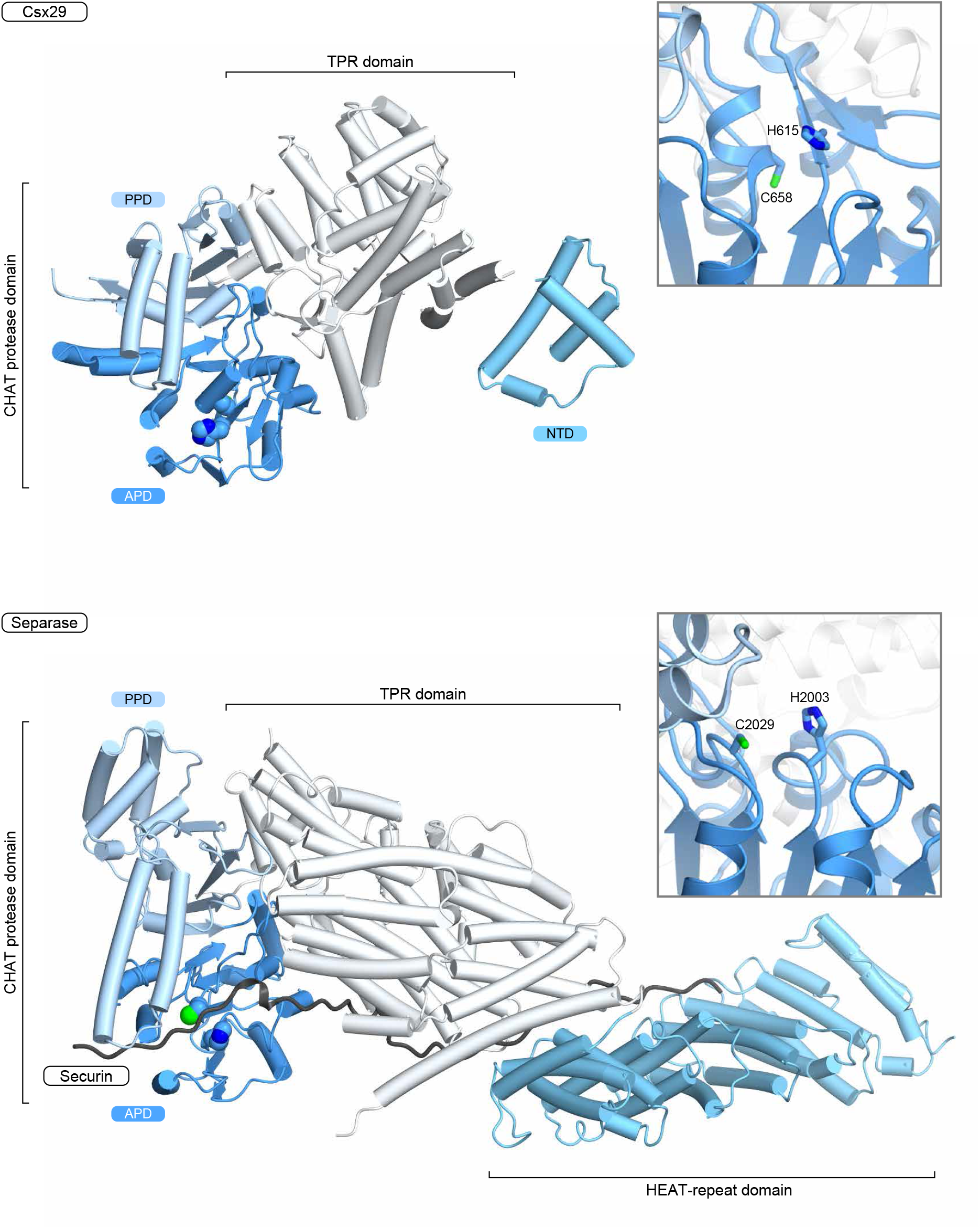
Structural comparison between Csx29 and human separase. Overall structures of Csx29 and human separase (PDB ID: 7NJ1). The catalytic residues are depicted as space-filling models. Securin (separase inhibitor) is colored gray. The close-up views of the protease active sites are shown in insets.

**fig. S6.**
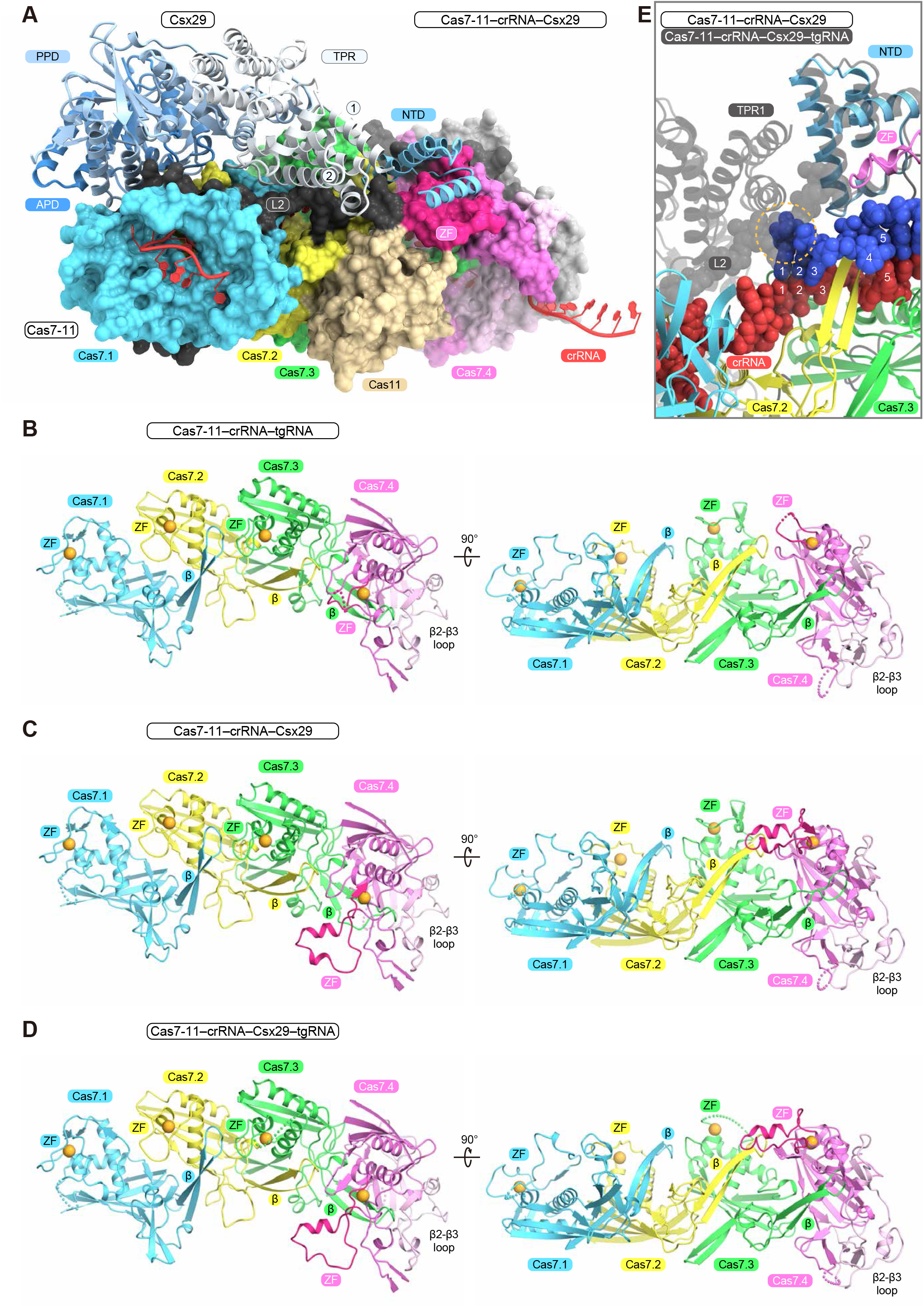
Interaction between Cas7-11 and Csx29. (A) Interface between Cas7-11 and Csx29 in the Cas7-11–crRNA–Csx29 complex. Cas7-11 and Csx29 are shown as ribbon and surface representations, respectively. The INS and CTE domains of Cas7-11 are omitted for clarity. (B–D) Structures of the Cas7.1–Cas7.4 domains in Cas7-11–crRNA–tgRNA (PDB ID: 7WAH) (B), Cas7-11–crRNA–Csx29 (C), and Cas7-11–crRNA–Csx29–tgRNA (D). The bound zinc ions are shown as orange spheres. The α-helical insertion in the Cas7.4 ZF motif is highlighted by magenta. (E) Potential steric clash between Cas7-11/Csx29 and a bound target RNA. The Cas7-11–crRNA–Csx29 structure was superimposed onto the Cas7-11–crRNA–Csx29–tgRNA structure, and then only Csx29 and Cas7-11 L2 are shown in semi-transparent gray. A potential steric clash between Cas7-11 L2/Csx29 TPR1 and the bound target RNA is indicated by a dashed orange circle.

**fig. S7.**
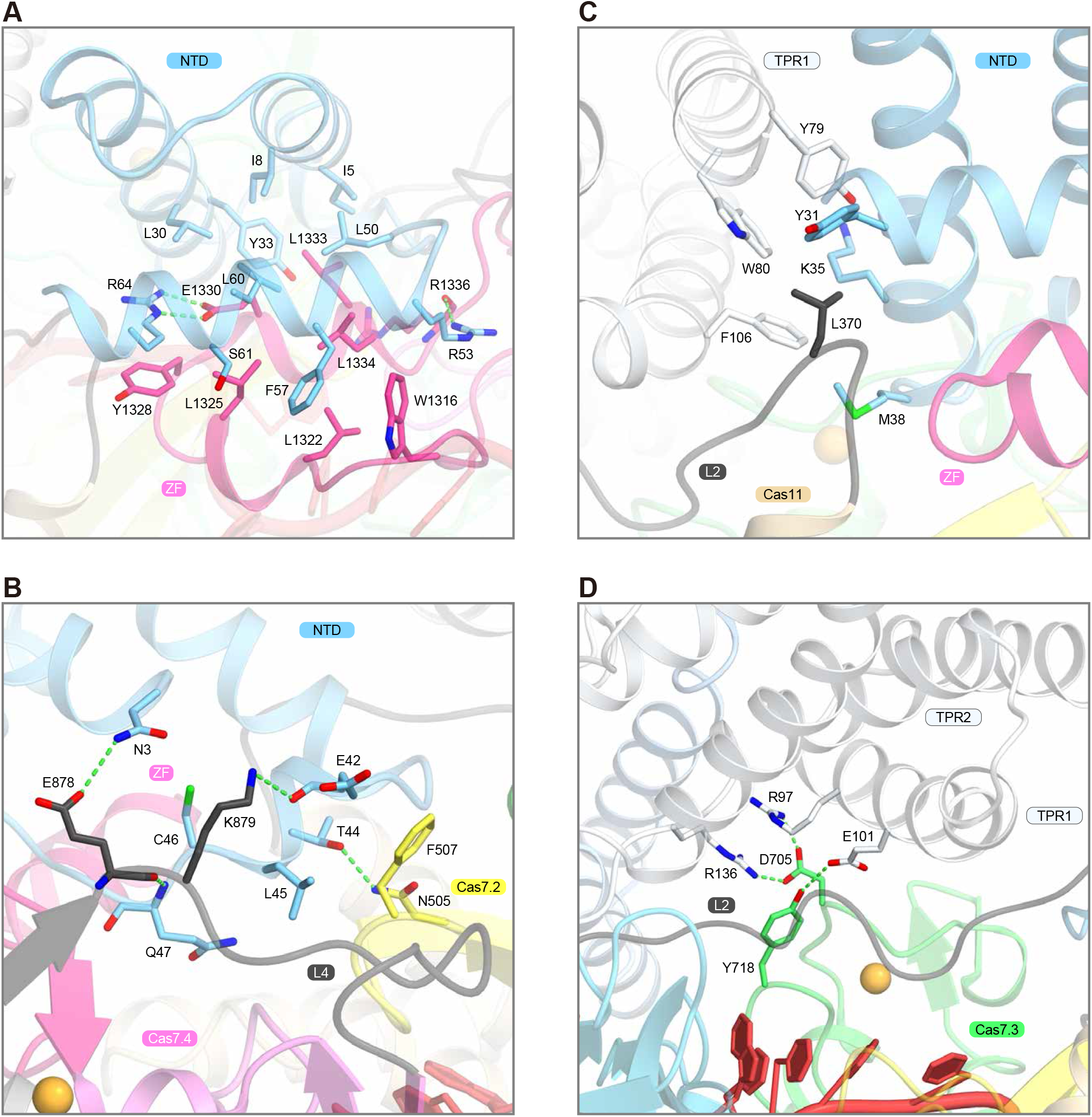
Interface between Cas7-11 and Csx29. (A) Interface between Cas7-11 Cas7.4 and Csx29 NTD. (B) Interface between Cas7-11 Cas7.3/L2 and Csx29 NTD. (C) Interface between Cas7-11 L2 and the Csx29 NTD/TPR. (D) Interface between Cas7-11 Cas7.3 and Csx29 TPR1/2.

**fig. S8.**
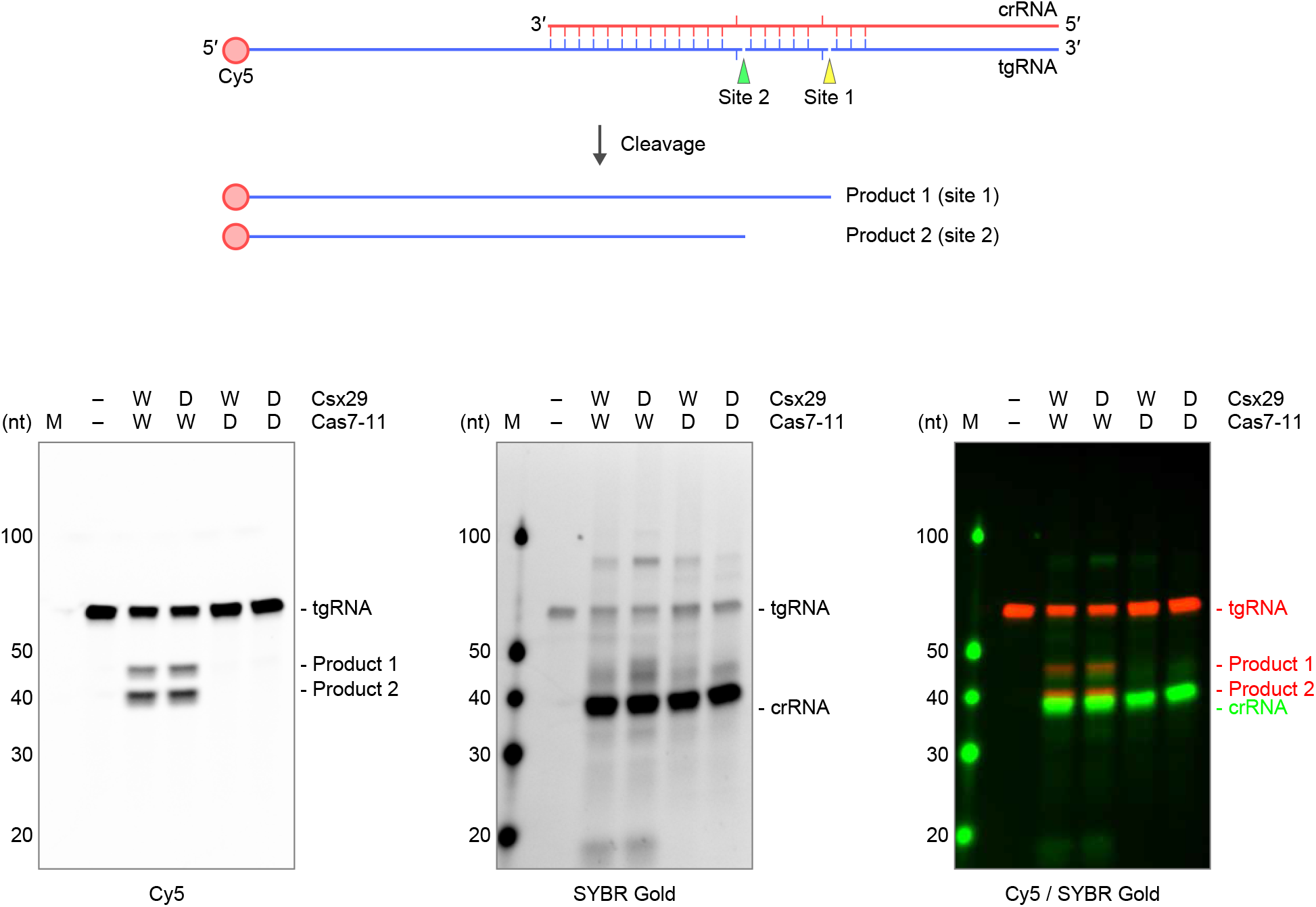
Target RNA cleavage by the Cas7-11–Csx29 complex. The Cas7-11–crRNA–Csx29 complex was incubated with a 5′-Cy5-labeled ssRNA target at 37°C for 10 min, and then analyzed by 15% TBE-urea PAGE. The gels were visualized, using either Cy5 or SYBR Gold fluorescence. The wild-type (W) and catalytically inactivated (D) versions of Cas7-11 and Csx29 were used.

**fig. S9.**
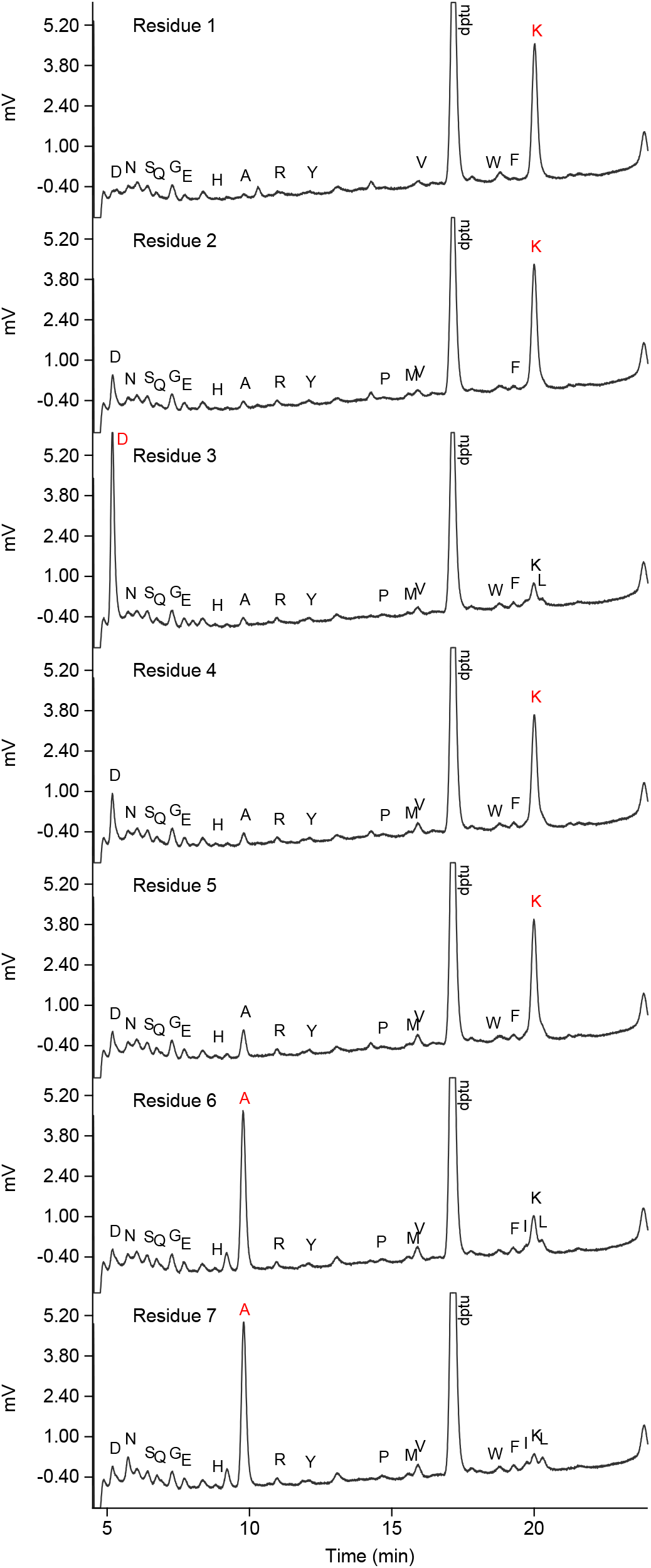
N-terminal analysis of Csx30. Elution profiles for N-terminal seven residues in the ~15 kDa Csx30 fragment (Csx30-2) were shown.

**fig. S10.**
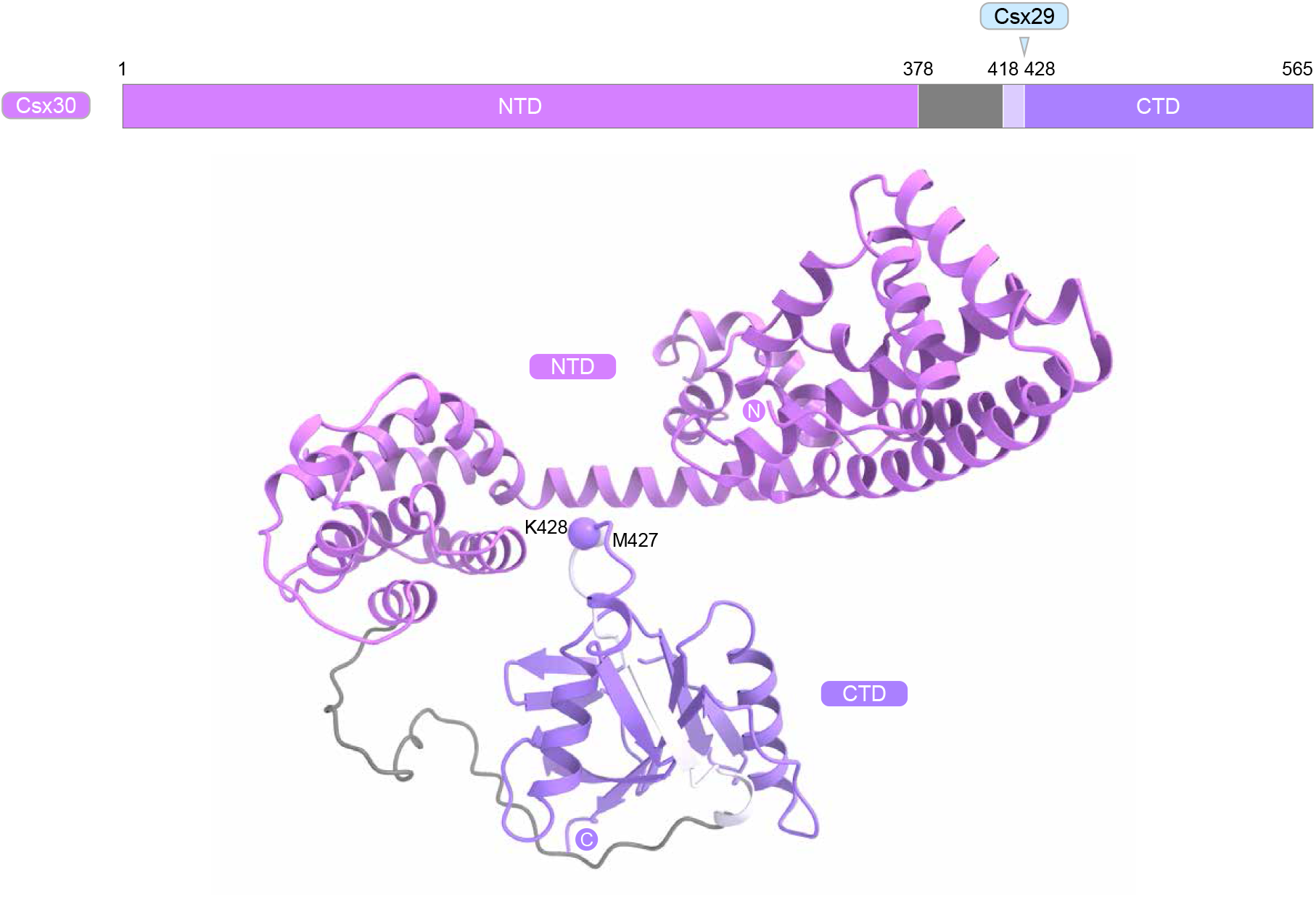
Predicted structures of Csx30. The Csx30 structure was predicted using AlphaFold2(*16*). The Cα atoms of M427 and K428 at the cleavage site are indicated by spheres.

**fig. S11.**
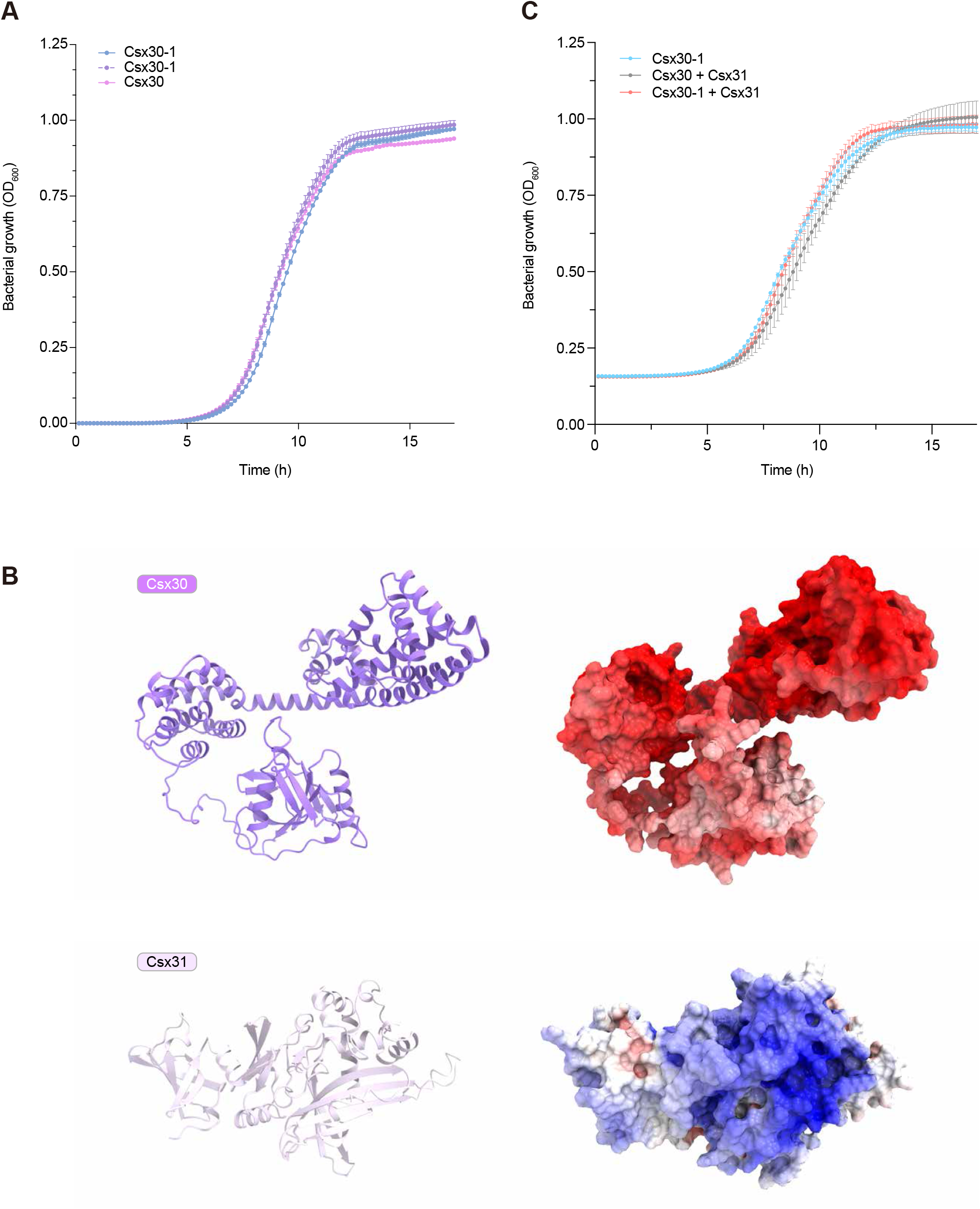
Effects of Csx30 and Csx31 on bacterial cell growth. (A) Growth curves of *E. coli* expressing the non-induced full-length Csx30, the N-terminal fragment (residues 1–427) of Csx30 (Csx30-1), or the C-terminal fragment (residues 428–565) of Csx30 (Csx30-2). These curves serve as non-induced controls for the curves in Fig. 4B. (B) Electrostatic surface potential of the Csx30 and Csx31 structures predicted using AlphaFold2(*16*). The predicted structures suggested that Csx30 and Csx31 have negatively and positively charged surfaces, respectively. (C) Growth curves of *E. coli* expressing non-induced Csx30-1, full-length Csx30 and Csx31, or Csx30-1 and Csx31. These curves serve as non-induced controls for the curves in Fig. 4D.

**fig. S12.**
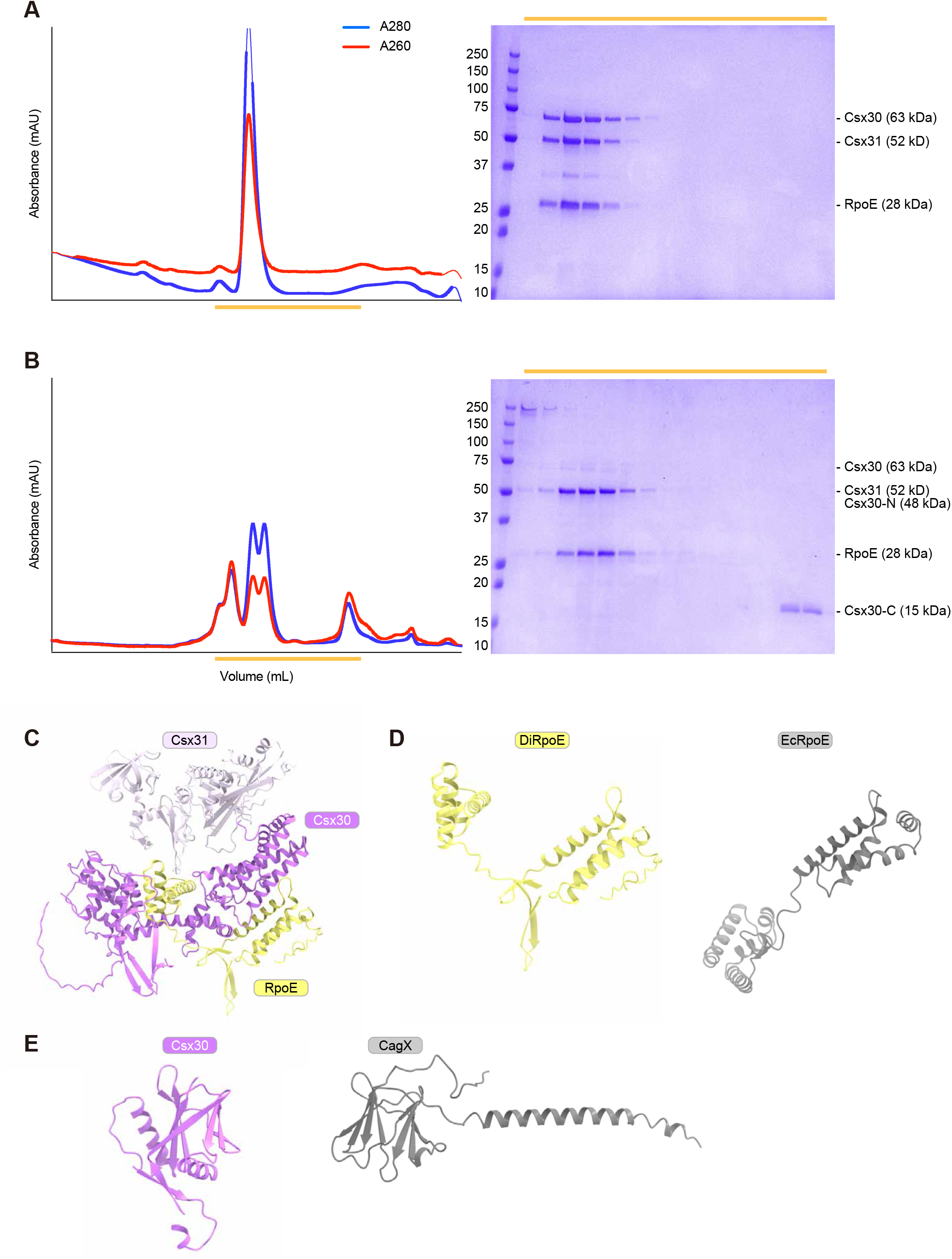
Interaction between Csx30, Csx31, and RpoE. (A and B) Elution profiles of the Csx30–Csx31–RpoE complex from a gel-filtration column. Csx30, Hig6-tagged Csx31, and Hig6-tagged RpoE were co-expressed in *E. coli*, and purified by Ni-NTA and HiLoad 16/600 Superdex 200. In (A), the Csx30–Csx31–RpoE complex was loaded onto Superdex 200 Increase. In (B), the Csx30–Csx31–RpoE complex was incubated with the Cas7-11–crRNA–Csx29–tgRNA complex, and then loaded onto Superdex 200 Increase. The fractions indicated by orange lines were analyzed by SDS-PAGE. The gels were stained with CBB. (C) Predicted structure of the Csx30–Csx31–RpoE complex. The structures of Csx30–Csx31 and Csx30–RpoE were predicted using AlphaFold2(*16*), and then they are superimposed based on the Csx30 NTDs. The Csx30 CTD in Csx30–RpoE is omitted for clarity. (D) Structural comparison of *D. ishimotonii* RpoE (model) and *E. coli* RpoE (PDB ID: 6JBQ). (E) Structural comparison of the Csx30 CTD (model) and CagX (PDB ID: 6OEG).

**fig. S13.**
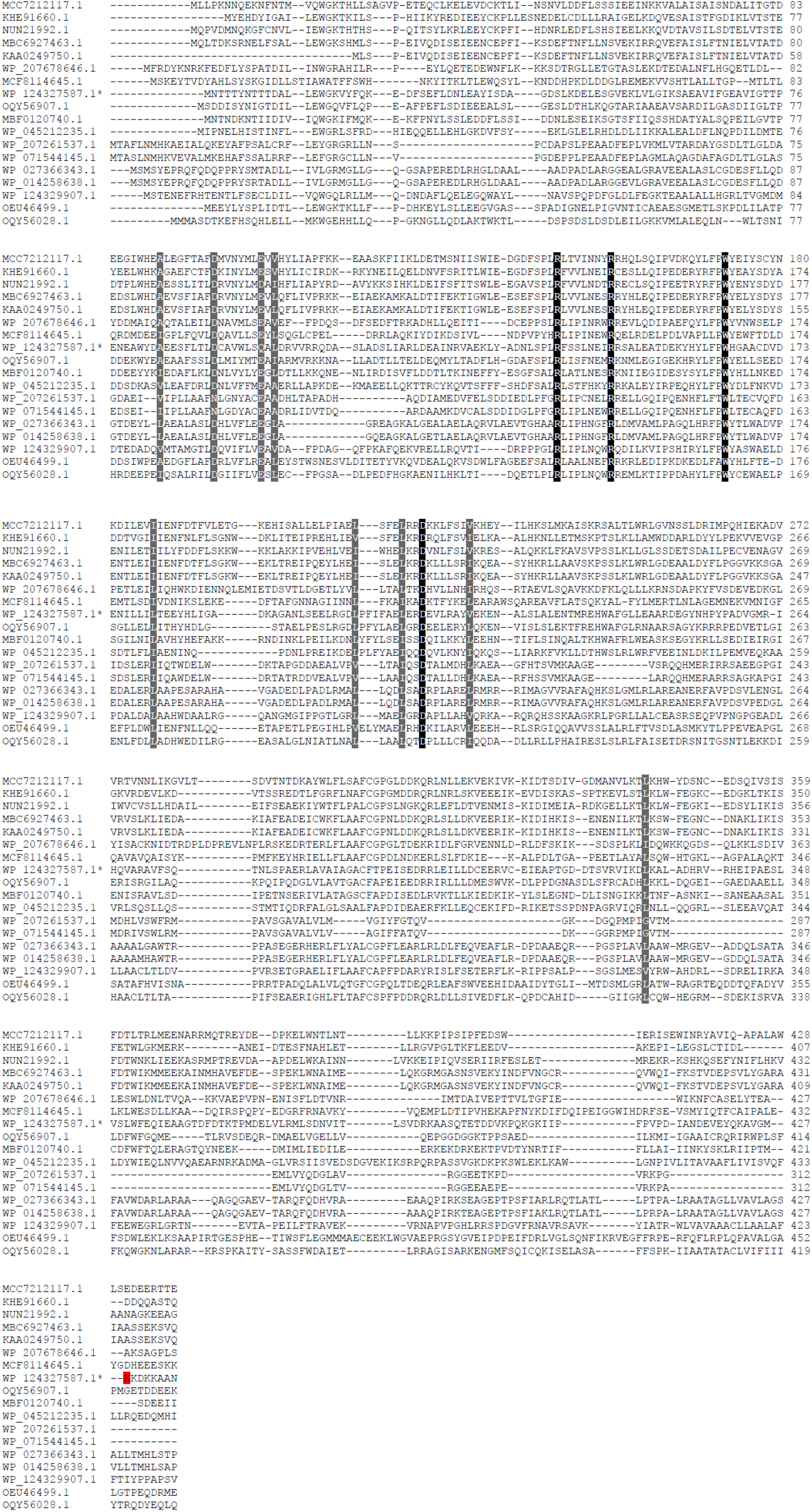
Multiple sequence alignment of the N-terminal domain of the Csx30 orthologs. Multiple alignment of the N-terminal domain of Csx30 family proteins was built using the Muscle5 program(*28*), and colored using Sequence Manipulation Suite (http://www.bioinformatics.org/sms2/color_align_cons.html). The Csx30 from *D. ishimotonii* is marked by an asterisk. Lysine residues, equivalent to K427 of *D. ishimotonii* Csx30 at the cleavage site by Csx29, is highlighted by red.

**fig. S14.**
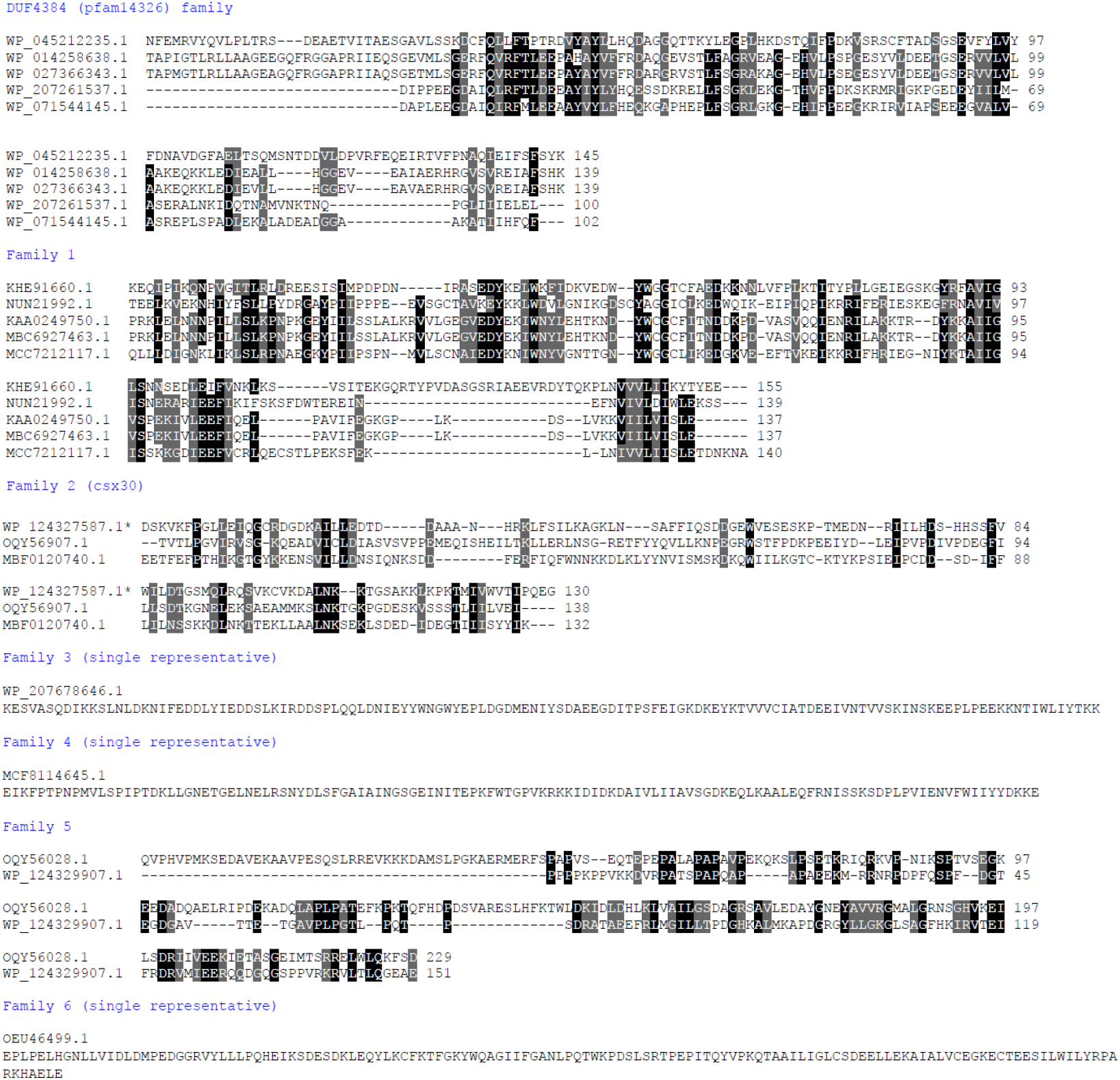
Multiple sequence alignment of the C-terminal domain of the Csx30 orthologs. Multiple alignment of the C-terminal domains of Csx30 family proteins were built using the Muscle5 program and colored using http://www.bioinformatics.org/sms2/color_align_cons.htmlserver. Three families are represented by a single sequence and are not therefore aligned.

