## Supplemental Text for "RNA-triggered protein cleavage and cell death by the RNA-guided type III-E CRISPR-Cas nuclease-protease complex"

### Table S1. Cryo-EM data collection, refinement, and validation statistics

| | Cas7-11–crRNA–Csx29 (EMDB-33695) (PDB 7Y7X) | Cas7-11–crRNA–Csx29–tgRNA (EMDB-33696) (PDB 7Y7Y) |
|---|---|---|
| **Data collection and processing** | | |
| Magnification | 105,000 | 105,000 |
| Voltage (kV) | 300 | 300 |
| Electron exposure ($e^-/Å^2$) | 48.5 | 48.4 |
| Defocus range (μm) | −0.8 to −2.0 | −0.8 to −2.0 |
| Pixel size (Å) | 0.83 | 0.83 |
| Symmetry imposed | *C*1 | *C*1 |
| Initial particle images (no.) | 3,538,872 | 4,275,699 |
| Final particle images (no.) | 692,038 | 291,630 |
| Map resolution (Å) | 2.48 | 2.77 |
| FSC threshold | 0.143 | 0.143 |
| **Refinement** | | |
| Model resolution (Å) | 2.65 | 2.95 |
| FSC threshold | 0.5 | 0.5 |
| Map sharpening *B* factor ($Å^2$) | 78.8 | 88.0 |

| | | |
|---|---|---|
| Model composition | | |
| Non-hydrogen atoms | 18,627 | 13,781 |
| Protein residues | 2192 | 1,545 |
| Nucleotide residues | 38 | 61 |
| Ligands | 4 | 4 |
| *B* factors (Å$^2$) | | |
| Protein | 47.94 | 39.14 |
| Nucleotide | 30.40 | 14.30 |
| Ligand | 75.26 | 74.44 |
| R.m.s. deviations | | |
| Bond lengths (Å) | 0.004 | 0.003 |
| Bond angles (°) | 0.626 | 0.603 |
| Validation | | |
| MolProbity score | 1.53 | 1.60 |
| Clashscore | 8.73 | 8.67 |
| Poor rotamers (%) | 1.20 | 1.48 |
| Ramachandran plot | | |
| Favored (%) | 98.61 | 98.49 |
| Allowed (%) | 1.39 | 1.51 |
| Disallowed (%) | 0.00 | 0.00 |

### Table S2. Csx29 and 30 plasmid maps

| **Name** | **Full Description** | **Benchling link** |
|---|---|---|
| Full-length wtCsx30 bacterial expression | Expression of Csx30 under arabinose-inducible promoter, kanamycin resistance | https://benchling.com/s/seq-Q8N6zZrjkadwhqQWdWFI?m=slm-UsJYgu75dA4PfEC2m68z |
| Csx30-1 bacterial expression | Expression of N-term Csx30 under arabinose-inducible promoter, kanamycin resistance | https://benchling.com/s/seq-dHVl08GZ3SPm5OM5K8F7?m=slm-NyNp8JdLh5on8sbrh9De |
| Csx30-2 bacterial expression | Expression of C-term Csx30 under arabinose-inducible promoter, kanamycin resistance | https://benchling.com/s/seq-XoNvTE5ZUo8cMxdGirXr?m=slm-vY9Hy9iGb63fP0EwJjZq |
| Full-length wtCsx31-Csx30 *cis* bacterial expression | Expression of wtCsx31-Csx30 with endogenous RBS under arabinose-inducible | https://benchling.com/s/seq-k9hlylchBnXsjbewceKR?m=slm-SKDnnzCdvqn6co4ymX1h |

| | | |
|---|---|---|
| | promoter, kanamycin resistance | |
| Full-length wtCsx31-Csx30-rpoE *cis* bacterial expression | Expression of wtCsx31-Csx30-rpoE with endogenous RBS under arabinose-inducible promoter, kanamycin resistance | https://benchling.com/s/seq-Shz0rSS3hWUXewSaPOFY?m=slm-SnkUthyZpjjiLotTvShW |
| Csx30-1–Csx31 bacterial expression | Expression of wtCsx31-N-term Csx30 with endogenous RBS under arabinose-inducible promoter, kanamycin resistance | https://benchling.com/s/seq-8nUKhBl0m1sSfEbfwgXX?m=slm-vvxiFmq3QD7zaLSnRy6F |
